# Alpha-BET: Functional labeling of envelope glycoproteins with single domain antibodies for in-virus single molecule imaging

**DOI:** 10.1101/2025.04.11.648405

**Authors:** David J. Willliamson, Cecilia Zaza, Irene Carlon-Andres, Alessia Gentili, James L. Daly, Harry Holmes, Joseph W. Thrush, Tobias Starling, Stuart Neil, Ray Owens, Michael H. Malim, Christopher Tynan, Sabrina Simoncelli, Sergi Padilla-Parra

## Abstract

We present Alpha-BET, a structure-guided strategy leveraging AlphaFold to identify optimal ALFA-tag insertion sites for minimally disruptive labeling of viral glycoproteins with high-affinity nanobodies. Applied to HIV-1 Env, SARS-CoV-2 S, and NiV G, Alpha-BET preserves structural integrity and function. For HIV-1 Env, we demonstrate super-resolution DNA-PAINT MINFLUX 3D imaging enabled by tag insertion, showcasing its power for visualizing native trimers in single virions and potential for broader applications in virus research.

## Main

Labeling viral envelope glycoproteins particularly Class I fusogens^1^, is crucial for understanding enveloped virus entry and rational vaccine design^2^. These viral glycoproteins undergo complex structural rearrangements that dictate viral entry and immune evasion. For instance, studies on the Nipah virus attachment (G) glycoprotein have revealed distinct pre-fusion and post-fusion states, highlighting key transitions necessary for viral entry^3^. Similarly, single-molecule Förster resonance energy transfer (smFRET) provided insights into the dynamic conformational changes of the SARS-CoV-2 spike (S) protein^4^. HIV-1 Env has also been extensively studied using smFRET^5^ and super-resolution STED microscopy^6^, demonstrating its intrinsic heterogeneity and the impact of antibody binding on its conformational landscape. The use of non-canonical amino acids (ncAAs) with amber codon suppression techniques has recently been used in smFRET studies of HIV-1 Env to achieve precise, site-specific tagging without disrupting the protein’s function^7^. By introducing an amber stop codon at key sites in the gp120 subunit, non-canonical amino acids (ncAAs) carrying bio-orthogonal chemical groups are incorporated, allowing for selective attachment of donor and acceptor fluorophores via click chemistry. Whilst perfect for smFRET studies, these labelling approaches are not suited for super-resolution fluorescence microscopy due to problems related to signal to noise, the effective number of viral particles labeled, and notably photobleaching^8^.

Single-domain antibodies (sdAbs), also known as ‘nanobodies’, are attractive candidates for super-resolution imaging labelling due to their small size, ease of production, and their potential for single-site reporter conjugation, e.g. to functionalize the nanobody with a dye or enzyme, etc. A high affinity nanobody-tag pair allows the precision and specificity of genetic labelling with the flexibility of reporter readouts for the same target. Many tagging approaches introduce the epitope sequence to the N- or C-terminus of the target protein. However, these sites may not be accessible to nanobody binding or may have important signaling functions. Internal labelling is more complex as it requires positioning the tag so as not to disturb the functionality of either the target protein or the tag.

To address these challenges, we present Alpha-BET (AlphaFold2^9^ assisted Bio-Enhanced Tagging) (Fig. 1a), a structure-guided strategy leveraging the predictive capabilities of AlphaFold2 to identify optimal insertion sites for the ALFA-tag^10^, a small, rationally designed synthetic epitope tag optimized for advanced fluorescence microscopy. Alpha-BET was successfully applied and validated across diverse viral envelope glycoproteins, including HIV-1 Env, SARS-CoV-2 S protein, and NiV G. The workflow consists of five key steps (Fig. 1a): A. Producing of a list of tagging proposals, B. Generation of structural models with AlphaFold, C. Examination of structural details: malformed tags and changes to protein conformation, D. Alignment and/or docking checks for likely nanobody engagement and potential steric clashes and E. Produce, test and validate optimal candidates.

**Figure 1.**
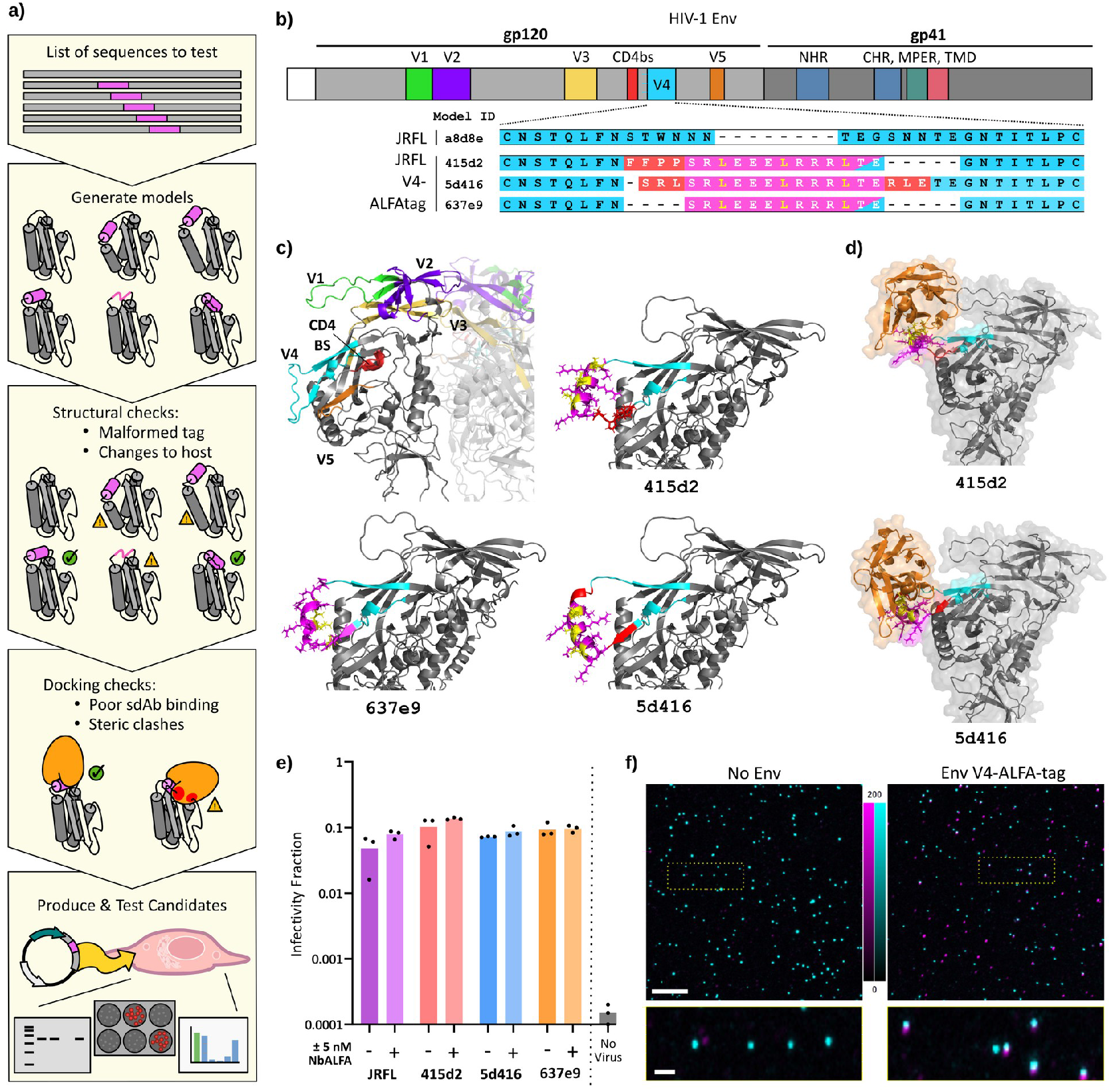
A general approach to screening tag inserts using AlphaFold. **(a)** A schematic of the major steps involved: compiling a list of sequences with different tag-insertion proposals, these are used to generate models in, e.g. AlphaFold, and inspected for structural problems, tag presentation, and nanobody docking, with favorable candidate passing to the bench for cloning, expression, and validation. **(b)** Major features of HIV-1 Env, and the context of three ALFA-tag labeling candidates 415d2, 5d416, and 637e9. **(c)** Structural model of JR-FL, with variable loops highlighted and AlphaFold2 models of the three candidate structures; key regions are the V4 loop (cyan), ALFA-tag (magenta), binding-pocked leucine residues (yellow), and padding residues (red). **(d)** NbALFA (orange) aligned to the ALFA-tag (magenta), **(e)** functional test by infectivity of ALFA-tag labelled Env with NbALFA in the with or without 5 nm NbALFA, and **(f)** functional test by imaging of ALFA-tag labelled Env (magenta) in viral particles (cyan).

In the case of the HIV-1 JR-FL Env, a representative CCR5-using envelope from a primary isolate, a collection of 26 tagging proposals were created wherein the ALFA-tag was inserted within the fourth variable (V4) loop (Fig. 1b, Supplementary Table 1). For each tagging proposal, AlphaFold2 generated five structural-prediction models. The crystal structure of the ALFA-tag bound to its nanobody (PDB ID: 6I2G) shows the tag as a straight alpha-helix, with three leucine residues exposed and facing the nanobody’s binding pocket (Fig. 1b and Supplementary Fig. 1-5). Therefore, the fitness of each tagging proposal was evaluated according to the shape and orientation of the ALFA-tag within the V4 loop. Tagging proposals were kept if their AlphaFold2 models displayed a straight alpha-helix which was oriented so that the binding-pocket leucines were solvent-accessible. In many cases, the ALFA-tag maintained an alpha-helical conformation but presented the binding-pocket leucines ‘inwards’, facing the bulk gp120 structure, which may impede accessibility and reduce the binding of the nanobody. Based on these assessments, for the HIV-1 Env JR-FL, two tagging proposals with favorable tag insertions (designated with Unique IDs 415d2 and 5d416) and one with an unfavorable insertion (UID 637e9) were selected for further study and validation.

Functional validation involved assessing protein expression, processing, and viral infectivity. Western blot analysis of HEK 293T lysates and HIV-1 particles decorated with either JR-FL WT or JR-FL-v4-ALFA-tag (UID 415d2) confirmed that the tagged precursor Env was processed properly to gp120 and gp41 subunits and incorporated into virions (Supplementary Fig. 6). To evaluate infectivity, viral samples expressing JR-FL WT, 415d2, 5d416, and 637e9 constructs were diluted (10-, 100-, and 1000-fold), with or without 5 nM anti-ALFA-tag nanobody (NbALFA) and exposed to HIV-1 sensitive TZM-bl reporter cells (Fig. 1e). The fraction of infected cells remained unchanged by ALFA-tag insertion in the V4 loop or the addition of nanobodies, indicating that these modifications did not impair viral envelope function. Cell-cell fusion assays also showed that the ALFA-tag insertion did not impede fusogenic activity (Supplementary Fig. 7). Additionally, fluorescence cross-correlation spectroscopy (FCCS) (Supplementary Fig. 8) and confocal co-localization (Fig 1f) confirmed the proper surface exposure of ALFAtag. FCCS analysis of HIV-1 viruses additionally labeled with the viral structural polyprotein Gag fused to EGFP and decorated with either JR-FL-v4-ALFA-tag (UID 415d2 or 637e9) revealed that only the 415d2 construct allowed successful docking of labeled NbALFA-ATTO 594, confirming proper ALFA-tag surface display. The labeling and validation of SARS-CoV-2 S Wuhan, vesicular stomatitis virus glycoprotein (VSV-G), and NiV G, also followed these steps (Supplementary Figs. 9-12) confirming the effectiveness of Alpha-BET to predict labelling efficiency and maintain functionality.

The successful implementation of Alpha-BET enables robust, function-preserving labeling of viral envelope glycoproteins for high-resolution microscopy. The ALFA-tag is particularly well-suited for DNA-PAINT^11^, a super-resolution method exploiting transient hybridization of DNA strands to achieve sub-5 nm localization precision—surpassing previous imaging approaches in viruses (Fig. 2a)^6,12^. We demonstrate this by imaging Env trimers on HIV-1 viruses decorated with the JR-FL V4-ALFA-tag (UID 415d2) Env. Env trimers were labeled with DNA-conjugated, ALFAtag-specific nanobodies, and DNA-PAINT imaging was performed using complementary ATTO 655- or Cy3B-labeled imager strands, for MINFLUX^13,14^ or total internal reflection (TIRF) excitation, respectively. HIV-1 particles were distinguishable from extracellular vesicles, which co-purify with viral particles, as those particles which also incorporated Gag fused to either EGFP or mTurquoise2. Three-dimensional DNA-PAINT MINFLUX (Fig. 2b) images of individual viral particles were obtained with an isotropic single-molecule localization precision of ∼ 5 nm, as determined by the spread of the estimated coordinates derived from the emission trace of single molecule events (Supplementary Fig. 13). Zoomed-in images revealed tightly clustered Env trimers, consistent with previous STED observations but now visualized with 4-fold improved resolution^15^. Under TIRF, two-dimension^16^ Cramér-Rao lower bound (CRLB) and nearest-neighbor-based metric (NeNa)^15^ precision of 3.6 nm and 4.9 nm, respectively (Fig. 2c and Supplementary Fig. 14). This NeNa value represents a resolution of 11.5 nm which sets a new benchmark in HIV imaging, enabling for the first time the unambiguous visualization of individual Env trimers (Fig. 2d). Each Env trimer comprises three gp120 subunits forming a triangular structure. Cryo-electron tomography studies^17^ reported trimer widths of approximately 13 nm at the widest point, narrowing to approximately 10 nm at the base. Supplementary Fig. 15 presents simulated trimer structures with varying inter-subunit distances, generated at the same resolution as our 2D DNA-PAINT experiments. These simulations suggest that the experimentally observed trimeric structures likely measure less than 10 nm in size. These dimensions might indeed correspond with the long sought after unresolved HIV-1 native closed Env conformation^18^. In conclusion, Alpha-BET enables structurally informed, site-specific tagging compatible with advanced imaging workflows. The resulting enhancement in labeling precision and functionality paves the way for dissecting protein organization at the nanoscale—not only in viral systems but across a broad range of membrane-associated protein complexes.

**Figure 2:**
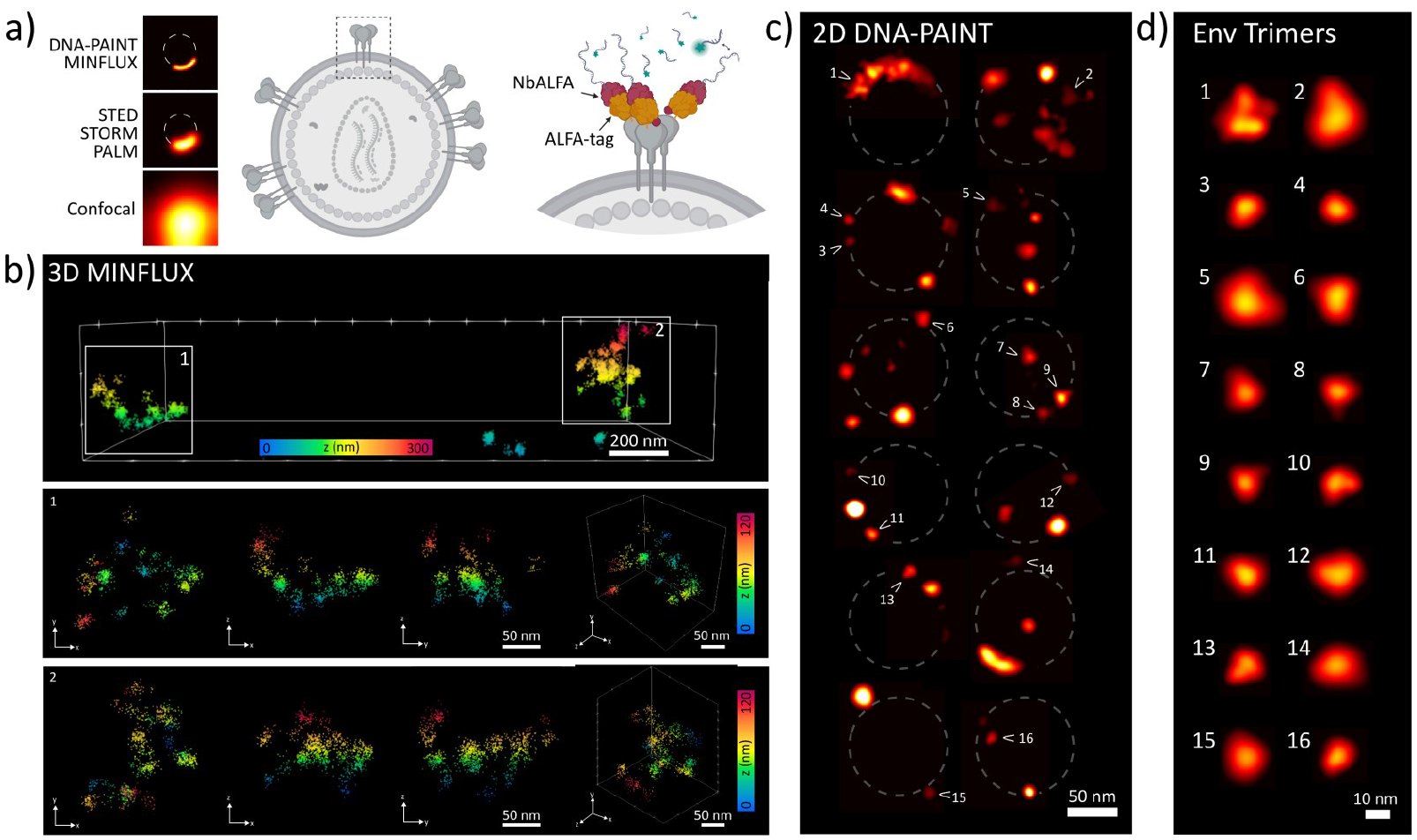
Applications to super-resolution imaging of viral particles. **(a)** Schematic of the resolution capabilities of various fluorescence microscopy techniques for visualizing Env clusters on a viral particle. **(b)** 3D DNA-PAINT MINFLUX imaging of individual viral particles, with two zoomed-in examples shown from different orientations. **(c)** Representative 2D DNA-PAINT images, obtained under TIRF illumination, of single viral particles; arrows indicate individual Env trimers. **(d)** Magnified view of the highlighted Env trimers shown in panel (c).

## Methods

### Cell Culture

All cells were grown in a humidified 5% CO_2_ atmosphere at 37°C. Cell culture media did not contain antibiotics.

Producer cells, Lenti-X 293T (Takara Bio, 632180), ACE2-expressing HEK 293T^+ACE2^, and Lenti-X 293TΔEFNB2 cells were maintained in phenol-red-free DMEM/F12 media (Thermo, 21041033) supplemented with 10% fetal bovine serum (FBS, Thermo).

Infectivity reporter cells, TZM-bl (NIH AIDS Research & Reference Reagent Program) and ACE2-expressing BEAS-2B^+ACE2^ cells, were maintained in DMEM (Thermo, 41965062) supplemented with 10% FBS.

A549^+ACE2^ were maintained in Kaighn’s modification of Ham’s F12 medium (Thermo, 21127022) supplemented with 10% FBS.

Lenti-X 293TΔEFNB2 were generated by CRISPR/Cas9 deletion^19, 20^ of the EFNB2 gene. Briefly, four gRNA target sites were identified in the vicinity of the EFNB2 start codon within the human genome (homo_sapiens, Ensembl release 112, chromosome 13:106489745-106535662). Oligo pairs were synthesized, annealed, and cloned into PX462 V2.0 plasmid (Addgene, #62987). Lenti-X 293T cells were co-transfected with all four plasmids and four days later subjected to selection with puromycin (Sigma). After three days, surviving cells were pooled, expanded, then subjected to NiV pseudoparticles to deplete the population of EFNB2-expressing cells through cell-cell fusion. The surviving cells were expanded and maintained for later use in cell-cell fusion experiments.

### Plasmid DNAs

Plasmids pR8ΔEnv (containing the HIV-1 genome with a deletion in Env), pcRev (expressing HIV-1 Rev) were kindly provided by Dr Greg Melikyan (Emory University, Atlanta, GA, USA). The pCAGGS plasmid containing JR-FL Env was a kind gift from Dr James Binley (Torrey Pines Institute for Molecular Studies, USA).

ALFA-tag sequence (SRLEEELRRRLTE) was introduced via a set of two PCR reactions meeting within the ALFA-tag insertion site. The ‘left’ PCR was performed with a CAG forward sequencing primer and a reverse primer with sequence homology to the V4 loop (as dictated by the insertion proposal) and half of the desired insertion sequence. The ‘right’ PCR was performed with a forward primer containing the other half of the desired insert sequence and homology to the remaining V4 loop, as dictated by the insertion proposal, and a rabbit beta-globin polyA reverse sequencing primer. The ‘left’ PCR product was digested with EcoRI (New England Biolabs) and the ‘right’ PCR product with XhoI (New England Biolabs). The ‘right’ fragment was phosphorylated using T4 Polynucleotide Kinase (New England Biolabs). The digested fragments were ligated into pCAGGS-JR-FL (cut with EcoRI and XhoI to remove the wildtype portion) using T4 Ligase (New England Biolabs), transformed into competent bacteria (NEB STABLE, New England Biolabs), and colonies screened for correctly oriented, tagged, insert.

## Oligonucleotides

### JRFL ALFA-tag

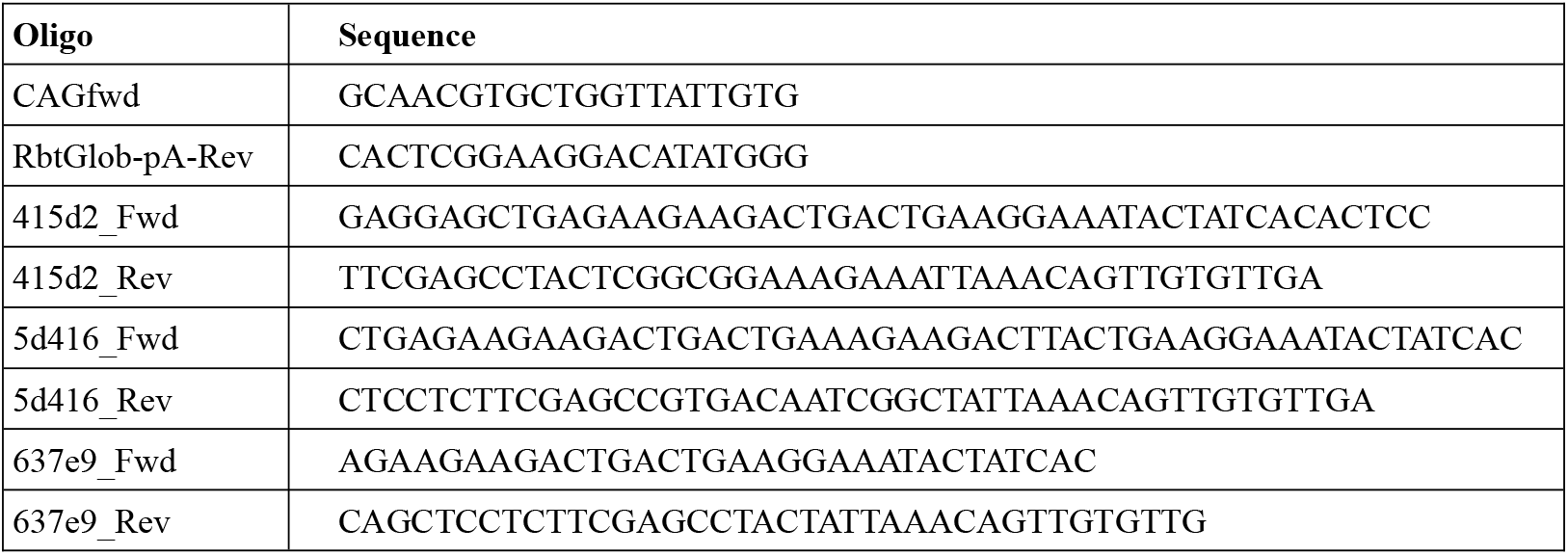

### EFNB2 gRNA

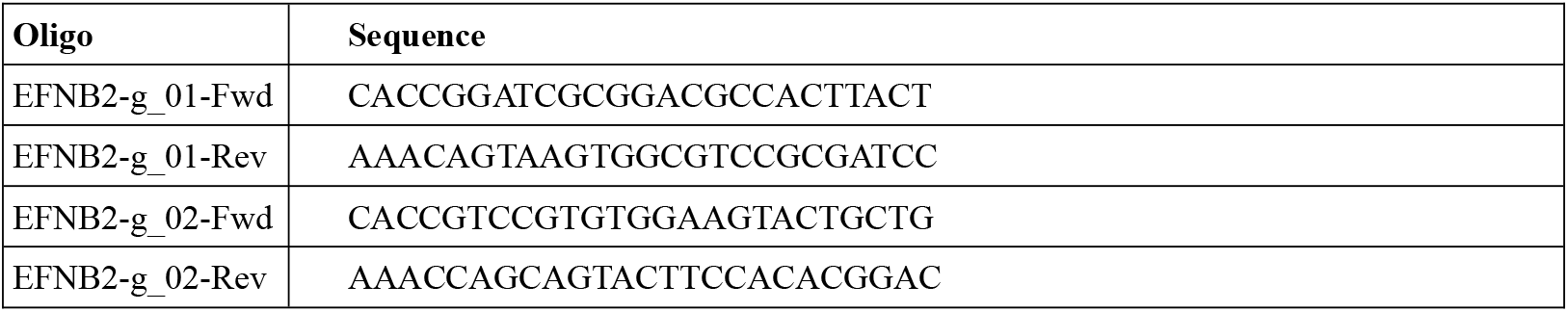
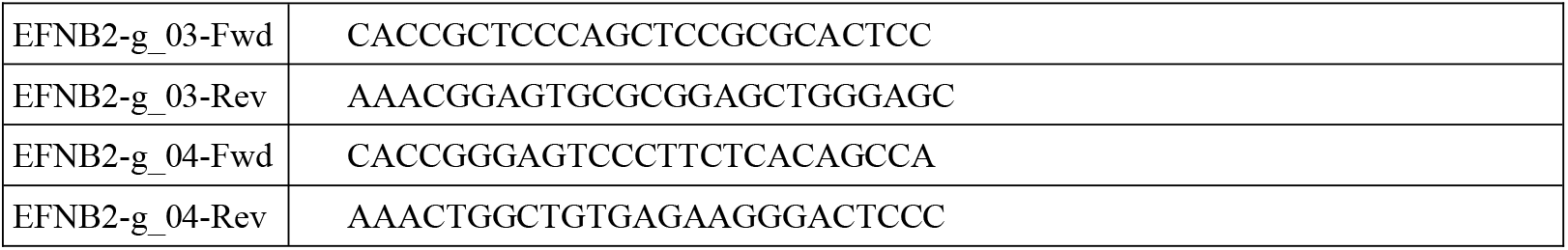

### sdAb labeling

Purified anti-ALFA-tag sdAbs (N1505, Nano-tag) bearing a single engineered cysteine were freshly reduced by adding 15 mM TCEP for 10 min on ice. PD-10 desalting columns (GE Healthcare) were used to exchange the buffer for maleimide labeling buffer (100 mM potassium phosphate pH 6.4, 150 mM NaCl, 1 mM EDTA, 250 mM sucrose, vacuum degassed and purged with nitrogen). For a standard labeling reaction, 10 nmoles of nanobody (i.e. 75–150 µM; 67-133 µl) were rapidly mixed with 12 nmoles of Alexa Fluor 647 C2 Maleimide (Life Technologies) (from a 20 mM stock in DMF), neutralized to pH 7.5 with K_2_HPO_4_ and incubated for 1.5 h on ice. Free dye was separated from labeled nanobody by buffer exchange to maleimide labeling buffer on PD10 desalting columns. Quantitative labeling was quality controlled by calculating the degree of labeling, which defines the molar ratio of dye to protein, as well as by SDS–PAGE and Coomassie staining.

### Modeling

The first 502 amino acids of HIV-1 Env JR-FL (GenBank: AAB05604.1), comprising the gp120 portion of the protein, together with modified sequences containing the ALFAtag, were used as the input for AlphaFold2 v2.2.0. Likewise for trimeric models of ALFA-tagged SARS-CoV-2 Spike, VSV-G, and NiV G. Structural model predictions were generated using a maximum template date of 22 February 2022, full genetic database, and monomeric model-preset options. Five models were created for each input sequence, and all were subjected to AMBER relaxation to minimize sidechain energy and remove steric clashes. Models were ranked by their pLDDT score and the highest scoring model then used for further investigation.

Models were visualized using PyMol (v2.5.0, Open-Source Build; The PyMOL Molecular Graphics System, Version 3.0 Schrödinger, LLC.). ALFAtag models were aligned to JR-FL and checked for gross distortions in the vicinity of the v4 loop compared to the wildtype structure. The ALFA-tag from PDB ID 6I2G (bound to nanobody) was aligned, in Pymol, to the ALFA-tag in the v4 loop of JR-FL. PyMol script ‘*show_bumps*’ was used to reveal steric clashes between the ‘bound’ nanobody and the tagged Env.

### Virus Production

Producer cells were seeded at a density of 65 × 10^3^ cells per cm^2^ in phenol-red-free DMEM/F12 medium. The following day the medium was replaced with FluoroBrite DMEM (Gibco, A1896701) supplemented with 3% FBS. The cells were then transfected using GeneJuice (Merck, 70967) at a GeneJuice to DNA ratio of 3:1. The DNA mixture contained pR8ΔEnv, pcRev, Gag-FP-ΔEnv (where FP is a fluorescent protein, e.g. EGFP, mTurquoise2, or mAmetrine), and Env at a plasmid copy ratio of 1:2:0.1:2.

For bulk production, cells were plated in T-175 vented flasks and transfected in 24 ml FluoroBrite medium with a total DNA load of 12 μg, combined with 36 μl of GeneJuice in 900 μl OptiMEM (Gibco 11058-21). Virion-containing media were harvested 48 hours after transfection and cleared of cell debris by centrifugation (5 minutes at 2000 × *g*) and passing the supernatant slowly through a 0.45 μm syringe filter. Filtered supernatant was then concentrated directly.

### Virus Concentration

Virions were cleaned and concentrated by low-speed centrifugation through a sucrose cushion^21, 22^. First, the volume of the medium after harvesting and initial clarification (approximately 24 ml from a T-175 producer flask) was adjusted to 28 ml with sucrose-free buffer (50 mM Tris-HCl, pH 7.4, 100 mM NaCl, 0.5 mM ethylenediaminetetraacetic acid (EDTA)). This was then underlaid with sucrose-cushion buffer (50 mM Tris-HCl, pH 7.4, 100 mM NaCl, 0.5 mM EDTA, and 292 mM sucrose (10 % w/v)) at a 4:1 (v/v) ratio and centrifuged at 10,000 × *g* for 3-4 hours, with minimal acceleration and deceleration profiles, at 4°C. After centrifugation the supernatant was carefully removed and the tubes inverted for 2 minutes to drain any residual supernatant. The pellet, containing virions, was gently overlaid with PBS (200-250 μl per pellet from a T-175 producer flask) and incubated for 30 minutes at 4°C. The pellet was then resuspended by very gentle mixuration in a pipette tip, dispensed into 20 μl aliquots and stored at -80°C. For experimental use, virions were thawed on ice and then used directly in sample preparation.

### Virus Titration

Virus infectivity and titers were determined using the TZM-bl cell line^23^, a line derived from HeLa and engineered to be permissive to HIV infection through the stable expression of CD4, CCR5, and endogenous expression of CXCR4. This line also contains stable inserts expressing, under the control of the HIV-1 long terminal repeat (LTR), β-galactosidase and luciferase as indicators of cell infection. Briefly, TZM-bl cells were plated at 60 × 10^3^ cells per cm^2^. The following day, aliquots of virus suspension were diluted and added to the reporter cells. After 48 hours, the cells were washed with PBS and fixed in 2% (v/v) paraformaldehyde (PFA) in PBS for ten minutes at 37°C. Fixed cells were then washed with PBS and incubated with X-Gal solution (5 mM K_3_[Fe(CN)]_6_ (potassium ferricyanide), 5 mM K_4_[Fe(CN)]_6_ (potassium ferrocyanide), 2 mM MgCl_2_, 1 mg/ml X-Gal (5-bromo-4-chloro-3-indolyl-β-D-galactoside, Thermo 15520034), in PBS) at 37°C for at least 2 hours. Cells were then washed with PBS and images acquired. The proportion of infected cells was calculated as the ratio of blue-stained (infected, β-galactosidase expressing) cells to the total number of cells originally plated, per unit area.

For luciferase-based infection assays, HEK 293T cells were transfected with p8.91, pCSLW and either JR-FL WT or JR-FL-v4-ALFA (UID 415d2) to produce pseudotyped lentiviral particles. U87-MG-CD4 cells expressing either CCR5 or CXCR4 were seeded in 96-well plates the day before infection. The next day, culture media was replaced with a dilution series of either JR-FL WT or JR-FL-v4-ALFA pseudotyped viruses, with or without a 30-minute pre-incubation step with 5nM anti-ALFA sdAb. Cells were returned to the incubator for 48 hours, then luminescence was quantified using the SteadyGlo assay (Promega) and a Victor X3 Multilabel Reader (Perkin Elmer).

### Western Blot

To identify JR-FL incorporation into virions, HEK 293T cells were transfected in a 10 cm dish with p8.91, pCSLW and either JR-FL WT or JR-FL-v4-ALFA (UID 415d2) to produce pseudotyped lentiviral particles. After 48 hours, supernatant was removed from cells, then the producer cells were lysed in 1% Triton X-100 PBS with protease inhibitors. 1mL of harvested supernatant was centrifuged at 18,000 × *g* for 2 hours to concentrate viral particles, then the pellet lysed in 1% Triton-X100 PBS with protease inhibitors. NuPAGE 4–12% gradient Bis-Tris precast gels (Life Technologies, NPO336) were used for SDS-PAGE, followed by transfer onto a polyvinylidine fluoride membrane (Immobilon-FL membrane, pore size 0.45 μm; Millipore, IPFL00010). The membrane was blocked, and sequentially labelled with anti-GAPDH (Proteintech), anti-p24 (24-2), or anti-gp41 (10E8) primary antibodies, or with FluoTag®-X2 anti-ALFA conjugated to LICOR IRDye 680RD (NanoTag). For anti-ALFA immunoprecipitation, anti-ACE2 EPR4436 (abcam, ab108209) was used to detect ACE2. Primary antibodies were subsequently labelled with fluorescent secondary antibodies (Invitrogen), Fluorescence detected by scanning with a LI-COR Odyssey Fc imager and Image Studio analysis software (LI-COR Biosciences).

### Anti-ALFA Immunoprecipitation

To test the ability of ALFA^Nterm^-Spike to interact with ACE2, 20 × 10^6^ HEK 293T stably expressing human ACE2 were transfected with 15 µg of ALFA-GFP, ALFA-VSV-G or ALFA^Nterm^-Spike and polyethylenimine (PEI) at a 1:3 DNA:PEI ratio. 48 hours later, cells were lysed on ice in 0.5% (v/v) NP40, 50 mM Tris-HCl, 150 mM NaCl lysis buffer. Lysates were centrifuged at 18,000 × *g* to remove insoluble debris, then the supernatants were incubated with ALFA Selector ST beads (NanoTag, N1511) and rotated for 1 hour at 4°C. Nonspecific interactors were removed by two washing steps with 0.25% NP40, 50 mM Tris-HCl, 150 mM NaCl wash buffer, followed by a final wash with 50 mM Tris-HCl, 150 mM NaCl wash buffer. All buffers contained protease inhibitors (Pierce, A32963) and were kept ice cold throughout the protocol. After the final wash was removed, proteins were eluted in 2x LDS samle buffer with 10% β-mercaptoethanol, and beads were boiled at 95°C for 10 minutes.

### Cell-cell fusion assay

Cell-cell fusion assays were performed to test the fusogenicity of ALFA-labelled envelope glycoproteins. In case of HIV-1 Envs, TZM-bl stably expressing CD4 and CCR5 HIV-1 receptors were used as target cells, and Lenti-X 293T cells expressing the different Envs as effector cells. 3000 TZM-bl cells stably expressing H2B-mTurquoise2 as nuclear reporter, were seeded in 300uL of DMEM (+10% FBS) per well, in a µ-Slide 8 Well (IBIDI, Cat No 80806). In parallel, 30 × 10^3^ Lenti-X 293T cells were seeded in 300 μL of DMEM F-12 (+10% FBS) per well. Lenti-X 293T were co-transfected with pCMV-eGFP, as cytosolic reporter, and the different Env expressing construcs: HIV-1 Env JRFL WT, JRFL V4-ALFA-tag (UID 415d2 or 5d416) or empty pcDNA (300ng plasmid DNA per well, in a 2:1 cytosolic reporter to Env ratio). In case of Nipah G proteins, Lenti-X 293T cells endogenously expressing EFNB2 Nipah receptor and transfected with pCMV-H2B-mTurquoise2 (150ng DNA plasmid per well), as nuclear marker, were used as target cells. Lenti-X 293TΔEFNB2 cells co-transfected with pCMV-eGFP, as cytosolic reporter, NiV F and NiV G WT, NiV G AF5, NiV G AF4 or empty pcDNA (300ng plasmid DNA per well, in a 2:0.5:0.5 cytosolic reporter to NiV F and NiV G ratio) were used as effector cells. Transfections were performed using GeneJuice reagent, following the manufacturer’s instructions. 24h post-transfection effector cells were detached with 0.5% Trypsin and added onto target cells. Cells were mixed in a 5:1 ratio in case of Lenti-X 293T added onto TZM-bl cells and 1:1 ratio in case of Lenti-X 293T WT added onto Lenti-X 293TΔEFNB2 cells. 24h after target and effector cell mixing, syncytia formation was observed using 2-color excitation confocal microscopy.

### DNA-PAINT & MINFLUX sample preparation

Glass bottom 6 channel Ibidi µ-slides (IB-80607) were cleaned with KOH 1 M for 2 h at room temperature and washed with copious volumes of clean water. The glass surface was then air-dried and exposed to UV light for 30 minutes. Viral particles were diluted 1:10 in PBS with 2% PFA and fixed at room temperature for ten minutes, then further diluted 1:10 in PBS. Fixed viral particles were added to the glass surface for 30 minutes and then the slide was washed 3 × 5 minutes with PBS, following 30 min blocking with PBS containing 2% BSA, 0.2% Fish Skin Gelatin. 25 nM DNA coupled anti-ALFA-tag nanobody (MASSIVE-TAG-Q-ANTI-alfa with R1: 5’-TTTCCTCCTCCTCCTCCTCCT-3’) in BSA 5% in PBS for 1 hour at room temperature. Samples were washed 3 × 5 minutes with PBS before the addition of 90 nm gold nanoparticles (G-90-100, CytoDiagnostics) for 5 minutes and a final 3 × 5 minutes with PBS. For imaging, DNA imager solution (R1: AGGAGGA) with 3’ attached Cy3b fluorophore in C+ buffer (1× PBS, 1 mM EDTA (Invitrogen), 500 mM NaCl (Sigma) pH 7.4, 0.02% Tween (Sigma)) was used. For 2D-DNA-PAINT-TIRF imaging a concentration of 2.5 nM of imager was used and for DNA-PAINT MINFLUX 1 nM imager concentration was used.

### DNA PAINT MINFLUX acquisition

MINFLUX imaging was performed on an Abberior instruments 3D MINFLUX microscope (Abberior instruments). Details of the microscope and components are described in Schmidt et al., 2021^24^. Briefly, the microscope is built on a motorized inverted microscope body (IX83, Olympus) with a CoolLED pE-4000 (CoolLED) for epifluorescence illumination. The system is equipped with 405 nm, 485 nm and 642 nm laser lines, as well as a 980 nm IR laser and xyz piezo (Physik Instrumente PInano XYZ) for the active sample stabilization system. Images were acquired with a 100× 1.4 NA UPLSAPO100XO oil objective (Olympus). During acquisition, the pinhole was set to 0.8 AU. MINFLUX acquisitions employed the combined fluorescence photons detected by two APDs with far red (650 – 685 nm) and near infrared (685 – 720 nm) spectral windows. The 485-nm confocal excitation beam enabled the detection of GFP fluorescence from the Gag-GFP core of the virus particles.

For 3D MINFLUX, imaging was performed with the default 3D imaging sequence supplied with the system. In brief, the sequence defines parameters (summarized below) used in the iterative localizing process of MINFLUX, culminating in a final pair of lateral and axial iterations with a TCP (targeted coordinate pattern) diameter of 40 nm, and minimum photon thresholds of 100 and 50 photons, respectively. Optical drift during acquisitions was corrected using the MINFLUX Beamline Monitoring (MBM) system, whereby MINFLUX measurements of fiducial marker positions are regularly taken between binding events.

| Iteration | L (nm) | TCP | Photon Threshold | cfr check | Dwell time (s) | Pattern Repeats | Background threshold (Hz) | Power Factor |
| --- | --- | --- | --- | --- | --- | --- | --- | --- |
| 0 | 288 | Hexagon | 160 | - | 0.001 | 1 | 15000 | 1 |
| 1 | 1400 | z-line | 400 | - | 0.001 | 1 | 15000 | 1 |
| 2 | 288 | Square | 100 | - | 0.001 | 5 | 10000 | 1 |
| 3 | 288 | z-line2 | 50 | - | 0.001 | 5 | 10000 | 1 |
| 4 | 151 | Square | 67 | 0.9 | 0.001 | 5 | 10000 | 2 |
| 5 | 151 | z-line2 | 33 | - | 0.001 | 5 | 10000 | 2 |
| 6 | 76 | Square | 67 | 0.8 | 0.001 | 5 | 10000 | 4 |
| 7 | 76 | z-line2 | 33 | - | 0.001 | 5 | 10000 | 4 |
| 8 | 40 | Square | 100 | - | 0.001 | 5 | 10000 | 6 |
| 9 | 40 | z-line2 | 50 | - | 0.001 | 5 | 10000 | 6 |

### DNA-PAINT MINFLUX data processing

A factor of 0.85 is applied to z-values to correct for foreshortening similar to Schmidt et al 2021^24^. This factor was determined for our system using 3D data from test samples of a known shape (centrioles and nucleopores).

Traces smaller than 5 localizations were discarded, and the remaining traces were filtered by the effective detection frequency to remove localizations that may be generated by multiple emitters. Each localization with a frequency greater than 42000 was removed. To avoid instances where filtered traces begin in one emitter position and end in another, each continuous section of a filtered trace was tagged with a new ID while preserving the original MINFLUX determined ID.

A histogram of the standard deviations of the localizations in each trace was plotted to show the variation in trace precision. The median trace standard deviation for xy and z define the localization precision in each axis (Supplementary Fig. 14a).

The localizations for each trace were centered by subtracting the mean trace position from each localization. These centered localizations were used to produce 2D histograms of *σ*_xy_ and *σ*_xz_ values. Ellipses with axis radii equal to 1, 2 and 3 standard deviations were plotted as appropriate (Supplementary Fig. 14b).

Finally, traces were aggregated as described in Schmidt et al with an effective photon count limit of 200 photons.

### Total internal reflection fluorescence (TIRF) microscopy setup

TIRF microscopy was carried out on a custom built total internal reflection fluorescence microscope based on a Nikon Eclipse Ti-2 microscope equipped with a 100× 1.49 NA oil immersion TIRF objective and a Perfect Focus System. Samples were imaged under flat-top TIR illumination with a 560 nm laser (MPB Communications, 1 W) magnified with both a custom-built telescope and a variable beam expander, before passing through a beam shaper device (piShaper 6_6_VIS, AdlOptica) to transform the Gaussian profile of the beam into a collimated flat-top profile. Laser polarization was adjusted to circular using a polarizer followed by a quarter wave-plate. The beam was focused into the back focal plane of the microscope objective using a suitable lens, passed through an excitation filter (FF01-390/482/563/640-25, Semrock) and coupled into the objective using a beam splitter (Di03-R405/488/561/635, Semrock). Fluorescent light was spectrally filtered with an emission filter (FF01-446/523/600/677-25, Semrock) and imaged on a sCMOS camera (Hamamatsu, ORCA-Fusion BT) without further magnification, resulting in a final pixel size of 130 nm in the focal plane, after 2 × 2 binning.

### DNA-PAINT TIRF acquisition and analysis

First, an image with a 488 nm CW laser was taken for 200 ms and a power density of 0.9 kW/cm^2^ to localize the position of the Gag-mAmetrine center of the virus particles on the previously described custom built microscope. Subsequent images were taken as a sequence of 100,000 frames, using an integration time of 100 ms and a density power of 2.3 kW/cm^2^.

The raw fluorescence videos were processed for super-resolution reconstruction using the Picasso software package^11^. Drift correction was applied using redundant cross-correlation, utilizing gold particles as fiducial markers for cellular experiments. Typically, around 50 nanoparticles were detected within the field of view, providing reliable drift correction, particularly for lateral shifts at the focal plane (z = 0).

Image rendering was carried out using the Render module of Picasso. Each localization was represented as a Gaussian spot, with the spread corresponding to the individual localization precision. For each pixel, the intensities of overlapping Gaussian spots were summed to determine the pixel intensity, which was then visualized using an appropriate colormap. To differentiate viruses from nonspecific binding or endosomes, the rendered DNA-PAINT image was aligned with the diffraction-limited image of the Gag channel using the gold nanoparticles. Only structures with the Gag signal were analyzed.

### Fluorescence Cross-Correlation Spectroscopy

Fluorescence cross-correlation Spectroscopy (FCCS) was performed using the MicroTime200 (Picoquant) time resolved fluorescence microscope. The incubator chamber (DigitalPixel) was preheated to 37°C. The sample was excited using pulsed 440 nm (for HIV-1 virions labelled with Gag-mTurquoise2) and 595 nm (for single domain antibodies labeled with ATTO 594) diode lasers (LDH series Picoquant) with a repetition rate of 20 MHz. The laser beam was coupled to an Olympus IX73 inverted microscope and focused onto the sample by a 63× 1.2 NA water immersion objective lens (Olympus UPlanXApo). Emission passed through a quad-dichroic mirror specifically designed to reflect four installed laser excitation lines (at 440, 485, 594, and 635 nm, Chroma) and a 300 μm pinhole (ThorLabs). The remaining emission was separated at a 560LP dichroic (Chroma). The longer-wavelength emission was then passed through 600LP and 690/70BP (Chroma) before arriving at a SPAD (Picoquant). The shorter-wavelength emission was passed through a 525/50B filter (Chroma) before a PMA hybrid detector (Picoquant). Time correlated single photon counting (TCSPC) was performed using the Multiharp 150 (Picoquant). The autocorrelation and cross-correlation curves for single viruses, single domain antibodies and single viruses engaged with single domain nanobodies were recovered and analyzed with Symphotime software (Picoquant) employing a free-diffusion, three-dimensional model^25^.

### Confocal Microscopy

Fusogenic SARS-CoV-2 Spike activity was assessed using confocal microscopy. Permissive cells, either BEAS-2B^+ACE2^ or A549^+ACE2^, were grown on 13 mm diameter #1.5 glass coverslips and transfected with p8.91, pCSLW and either wildtype Spike, ALFA^Nterm^-Spike, or ALFA^SD1^-Spike to produce pseudotyped lentiviral particles. After 24 hours, cells were fixed in 4% paraformaldehyde, permeabilized in 0.1% Triton X-100, blocked in 1% bovine serum albumin, and labeled with DAPI, CellMask Deep Red actin tracking stain (Invitrogen A57248), and FluoTag-X2 anti-ALFA conjugated to ATTO 488 (NanoTag). Images were acquired using a Nikon AXR inverted confocal microscope, and figures were prepared in Fiji.

### 2D-DNA-PAINT simulations

The simulations of the trimers shown in Supplementary Fig. 16 were performed using the Simulate package of Picasso software^11^. Parameters such as kinetic rates, imager strand concentration, excitation power density, background level, camera integration time, pixel size, and total number of frames were set to match the experimental conditions, ensuring comparable localization precision and NeNA values.

## Additional Software

Data analysis, statistical comparisons, and graphing were performed with Prism (version 9.1, GraphPad Software).

## Contributions

Conceptualization: D.J.W., S.P.-P.; methodology and validation: D.J.W., C.Z., I.C.-A., J.D., H.H., T.S., A.G., C.T., S.N., R.O., M.M., S.S., and S.P.-P.; formal analyses: D.J.W., C.Z., S.S.; resources: R.O., S.M., M.M., S.S., and S.P-P.; data curation: D.J.W., C.Z., S.S., C.T., and S.P.-P.; writing & original draft preparation: S.P.-P., and D.J.W.; writing, review, and editing: D.J.W., and S.P.-P.; visualization: D.J.W., C.Z., S.S., and S.P.-P.; supervision: S.S. and S.P.-P.; project administration: S.P.-P.; funding acquisition: R.O., S.N., M.M., S.S., and S.P.-P.. All authors have read and agreed to the published version of the paper.

## Acknowledgements

We thank the Nikon Imaging Centre at Kings College London for help with the AXR confocal microscopy

## References

Kielian, M. & Rey, F. A. Virus membrane-fusion proteins: more than one way to make a hairpin. Nature Reviews Microbiology 4, 67–76 (2006). 10.1038/nrmicro1326

Walker, L. M. & Burton, D. R. Rational antibody-based HIV-1 vaccine design: current approaches and future directions. Curr Opin Immunol 22, 358–366 (2010). 10.1016/j.coi.2010.02.012

Wang, Z. et al. Architecture and antigenicity of the Nipah virus attachment glycoprotein. Science 375, 1373–1378 (2022). 10.1126/science.abm5561

Lu, M. et al. Real-Time Conformational Dynamics of SARS-CoV-2 Spikes on Virus Particles. Cell Host Microbe 28, 880-891.e888 (2020). 10.1016/j.chom.2020.11.001

Munro, J. B. et al. Conformational dynamics of single HIV-1 envelope trimers on the surface of native virions. Science 346, 759–763 (2014). 10.1126/science.1254426

Chojnacki, J. et al. Maturation-dependent HIV-1 surface protein redistribution revealed by fluorescence nanoscopy. Science 338, 524–528 (2012). 10.1126/science.1226359

Ao, Y. et al. Bioorthogonal click labeling of an amber-free HIV-1 provirus for in-virus single molecule imaging. Cell Chem Biol 31, 487-501.e487 (2024). 10.1016/j.chembiol.2023.12.017

Roy, R., Hohng, S. & Ha, T. A practical guide to single-molecule FRET. Nat Methods 5, 507–516 (2008). 10.1038/nmeth.1208

Jumper, J. et al. Highly accurate protein structure prediction with AlphaFold. Nature 596, 583–589 (2021). 10.1038/s41586-021-03819-2

Götzke, H. et al. The ALFA-tag is a highly versatile tool for nanobody-based bioscience applications. Nat Commun 10, 4403 (2019). 10.1038/s41467-019-12301-7

Schnitzbauer, J., Strauss, M. T., Schlichthaerle, T., Schueder, F. & Jungmann, R. Super-resolution microscopy with DNA-PAINT. Nature Protocols 12, 1198–1228 (2017). 10.1038/nprot.2017.024

Carlon-Andres, I., Malinauskas, T. & Padilla-Parra, S. Structure dynamics of HIV-1 Env trimers on native virions engaged with living T cells. Commun Biol 4, 1228 (2021). 10.1038/s42003-021-02658-1

Balzarotti, F. et al. Nanometer resolution imaging and tracking of fluorescent molecules with minimal photon fluxes. Science 355, 606–612 (2017). 10.1126/science.aak9913

Ostersehlt, L. M. et al. DNA-PAINT MINFLUX nanoscopy. Nat Methods 19, 1072–1075 (2022). 10.1038/s41592-022-01577-1

Chen, Y. C. et al. Super-Resolution Fluorescence Imaging Reveals That Serine Incorporator Protein 5 Inhibits Human Immunodeficiency Virus Fusion by Disrupting Envelope Glycoprotein Clusters. ACS Nano 14, 10929–10943 (2020). 10.1021/acsnano.0c02699

Endesfelder, U., Malkusch, S., Fricke, F. & Heilemann, M. A simple method to estimate the average localization precision of a single-molecule localization microscopy experiment. Histochem Cell Biol 141, 629–638 (2014). 10.1007/s00418-014-1192-3

Li, Z. et al. Subnanometer structures of HIV-1 envelope trimers on aldrithiol-2-inactivated virus particles. Nature Structural & Molecular Biology 27, 726–734 (2020). 10.1038/s41594-020-0452-2

Lu, M. et al. Associating HIV-1 envelope glycoprotein structures with states on the virus observed by smFRET. Nature 568, 415–419 (2019). 10.1038/s41586-019-1101-y

Sanjana N.E., Shalem O., & Zhang, F. Improved lentiviral vectors and genome-wide libraries for CRISPR screening. Nature Methods 11 783–784 (2014). 10.1038/nmeth.3047

Shalem O. et al. Genome-scale CRISPR-Cas9 knockout screening in human cells. Science 343 83–7 (2014). 10.1126/science.1247005

Jiang, W. et al. An optimized method for high-titer lentivirus preparations without ultracentrifugation. Scientific Reports 5 (2015). 10.1038/srep13875

Boroujeni, M. E. & Gardaneh, M. The Superiority of Sucrose Cushion Centrifugation to Ultrafiltration and PEGylation in Generating High-Titer Lentivirus Particles and Transducing Stem Cells with Enhanced Efficiency. Molecular Biotechnology 60 (2018). 10.1007/s12033-017-0044-5

Wei, X. et al. Emergence of Resistant Human Immunodeficiency Virus Type 1 in Patients Receiving Fusion Inhibitor (T-20) Monotherapy. Antimicrobial Agents and Chemotherapy 46, 1896–1905 (2002). 10.1128/aac.46.6.1896-1905.2002

Schmidt, R. et al. MINFLUX nanometer-scale 3D imaging and microsecond-range tracking on a common fluorescence microscope. Nature Communications 12, 1478 (2021). 10.1038/s41467-021-21652-z

Padilla-Parra, S., Audugé, N., Coppey-Moisan, M. & Tramier, M. Dual-color fluorescence lifetime correlation spectroscopy to quantify protein-protein interactions in live cell. Microsc Res Tech 74, 788–793 (2011). 10.1002/jemt.21015

