## Supplementary Information for "Alpha-BET: Functional labeling of envelope glycoproteins with single domain antibodies for in-virus single molecule imaging"

#### Supplementary Table 1 – List of ALFA-tag insertion proposals

| UID | V4 ... | Padding | ALFA-Tag | Padding | ... V4 |
|---|---|---|---|---|---|
| 0c6e4 | CNSTQLFNSTWNNN | | SRLEEELRRRLTE | | GNTITLPC |
| 0de2e | CNSTQLFN | PP | SRLEEELRRRLTE | | GNTITLPC |
| 16a67 | CNSTQLFNSTWNNN | | SRLEEELRRRLTE | | TEGSNNTEGNTITLPC |
| 181ca | CNSTQLFNSTW | A | SRLEEELRRRLTE | PP | GNTITLPC |
| 19532 | CNSTQLFNSTWNNN | P | SRLEEELRRRLTE | P | TEGSNNTEGNTITLPC |
| 1af0b | CNSTQLFNSTW | PPPA | SRLEEELRRRLTE | A | GNTITLPC |
| 22584 | CNSTQLFNSTWNNN | A | SRLEEELRRRLTE | | GNTITLPC |
| 3c53d | CNSTQLFN | P | SRLEEELRRRLTE | P | GNTITLPC |
| 415d2 * | CNSTQLFN | FFPP | SRLEEELRRRLTE | | GNTITLPC |
| 5c2d5 | CNSTQLFNSTW | P | SRLEEELRRRLTE | P | GNTITLPC |
| 5d416 * | CNSTQLFN | SRL | SRLEEELRRRLTE | RRL | TEGNTITLPC |
| 637e9 * | CNSTQLFN | | SRLEEELRRRLTE | | GNTITLPC |
| 6bbb1 | CNSTQLFNSTWNN | PPP | SRLEEELRRRLTE | A | GNTITLPC |
| 7455d | CNSTQLFNSTWN | PPP | SRLEEELRRRLTE | PP | GNTITLPC |
| 7c3c9 | CNSTQLFNSTWNN | PP | SRLEEELRRRLTE | P | GNTITLPC |
| a896a | CNSTQLFNSTWN | PP | SRLEEELRRRLTE | A | GNTITLPC |
| ae744 | CNSTQLFNSTW | P | SRLEEELRRRLTE | PP | GNTITLPC |
| bc3ad | CNSTQLFN | FFP | SRLEEELRRRLTE | PFF | GNTITLPC |
| c449b | CNSTQLFNSTWNNN | PP | SRLEEELRRRLTE | P | GNTITLPC |
| c7e56 | CNSTQLFNSTWNN | PA | SRLEEELRRRLTE | AP | TEGNTITLPC |
| cd7bd | CNSTQLFN | TQL | SRLEEELRRRLTE | TQL | GNTITLPC |
| d4935 | CNSTQLFNSTW | PPP | SRLEEELRRRLTE | A | GNTITLPC |
| edfad | CNSTQLFNSTWNNN | | SRLEEELRRRLTE | | GSNNTEGNTITLPC |
| f0018 | CNSTQLFNSTWNN | P | SRLEEELRRRLTE | P | GNTITLPC |
| f8943 | CNSTQLFNSTW | P | SRLEEELRRRLTE | | GNTITLPC |
| ffecd | CNSTQLFNSTWNN | FP | SRLEEELRRRLTE | P | GNTITLPC |

a8d8e (JR-FL WT) CNSTQLFNSTWNNNTEGSNNTEGNTITLPC

* Expressed and tested

TE = ALFA-tag & V4 homology

### Supplementary Figures

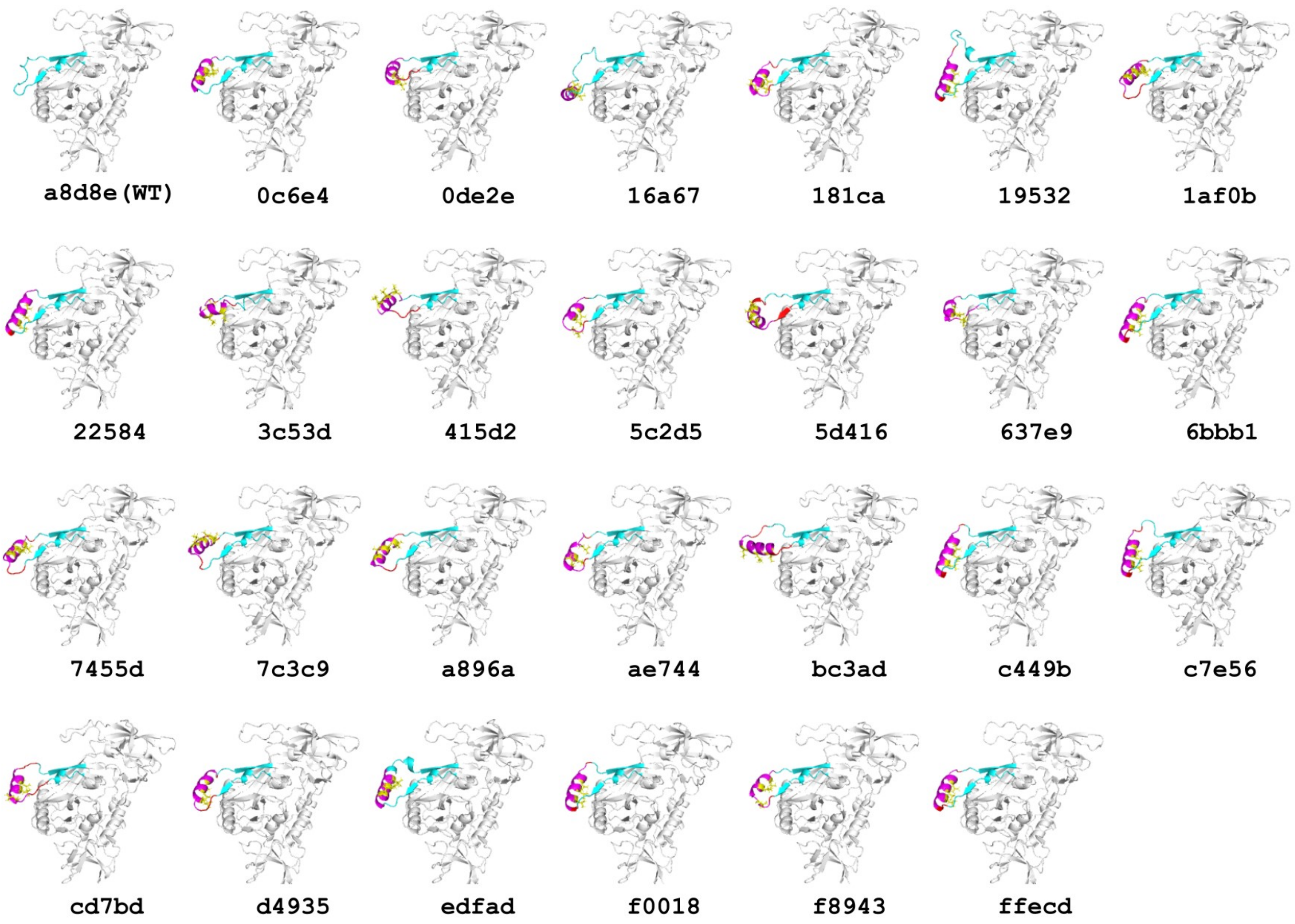


**Supplementary Figure 1. Overview of JR-FL wildtype (WT) and V4-ALFAtag labelled gp120 structures.** Alphafold2 structure predictions (and UID labels) of the JR-FL Env gp120 with ALFAtag (magenta) inserted at different positions within the variable loop 4 (cyan). Binding pocket leucines are shown in yellow. The highest ranked structure model for each input sequence is shown.

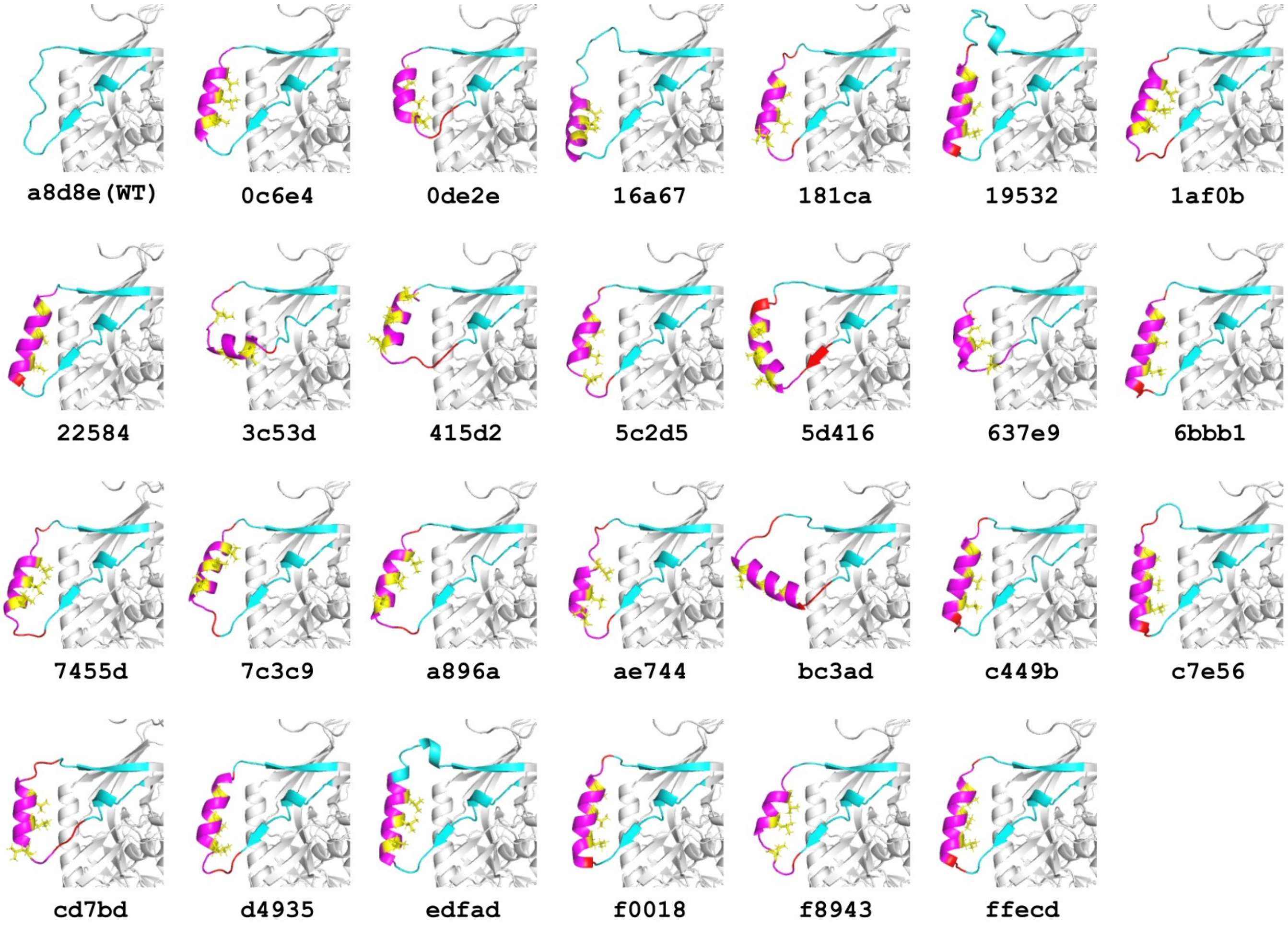


**Supplementary Figure 2. Zoomed, rotated view (of SI Fig. 1) showing detail of the V4 loop of the WT and labelled gp120 structures.** Alphafold2 structure predictions (and UID labels) of the JR-FL Env gp120 with ALFAtag (magenta) inserted at different positions within the variable loop 4 (cyan). Binding pocket leucines are shown in yellow. The highest ranked structure model for each input sequence is shown.

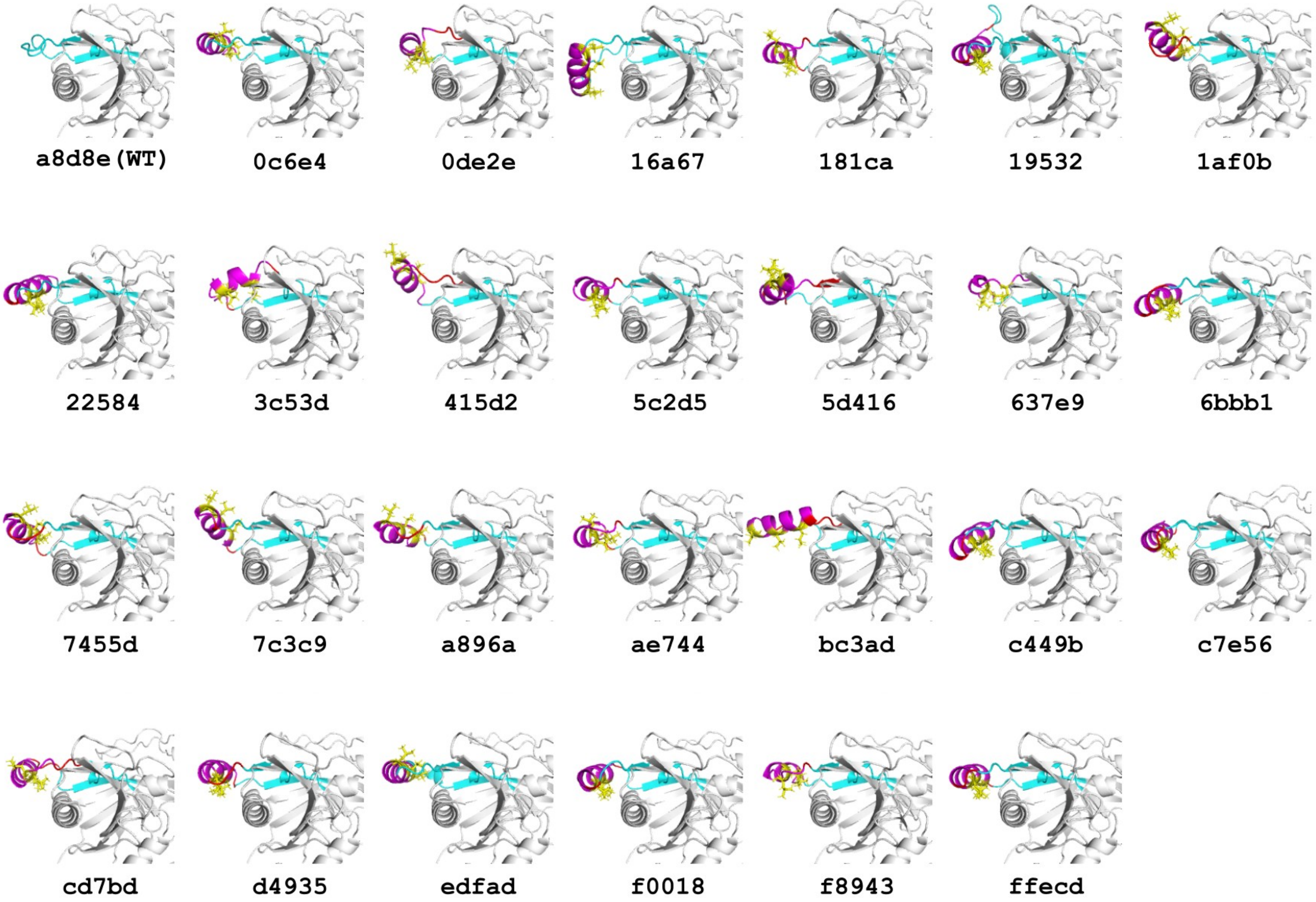


**Supplementary Figure 3. Zoomed, rotated view (of SI Fig. 1) showing detail of the V4 loop of the WT and labelled gp120 structures.** Alphafold2 structure predictions (and UID labels) of the JR-FL Env gp120 with ALFAtag (magenta) inserted at different positions within the variable loop 4 (cyan). Binding pocket leucines are shown in yellow. The highest ranked structure model for each input sequence is shown.

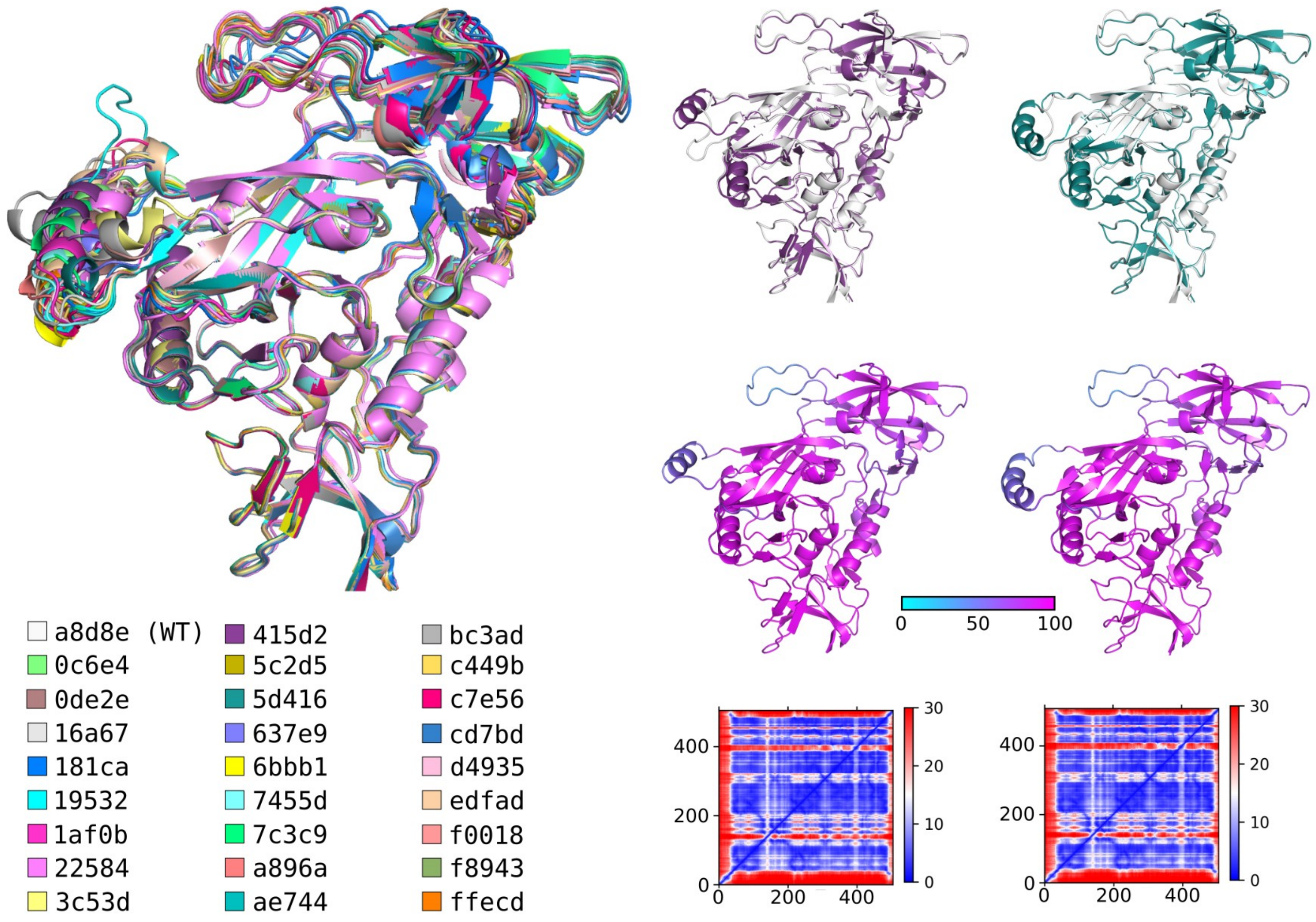


**Supplementary Figure 4. Alignment of tagging proposals to wildtype JR-FL.** a) The highest-ranked model of each tagging proposal aligned to the wildtype model, b) model UIDs 415d2 (purple) and 5d416 (teal) aligned to JR-FL WT (grey), c) models 415d2 and 5d416 colored according to predicted Local Distance Difference Test (pLDDT) and d) models' Predicted Aligned Error (PAE) heatmaps.

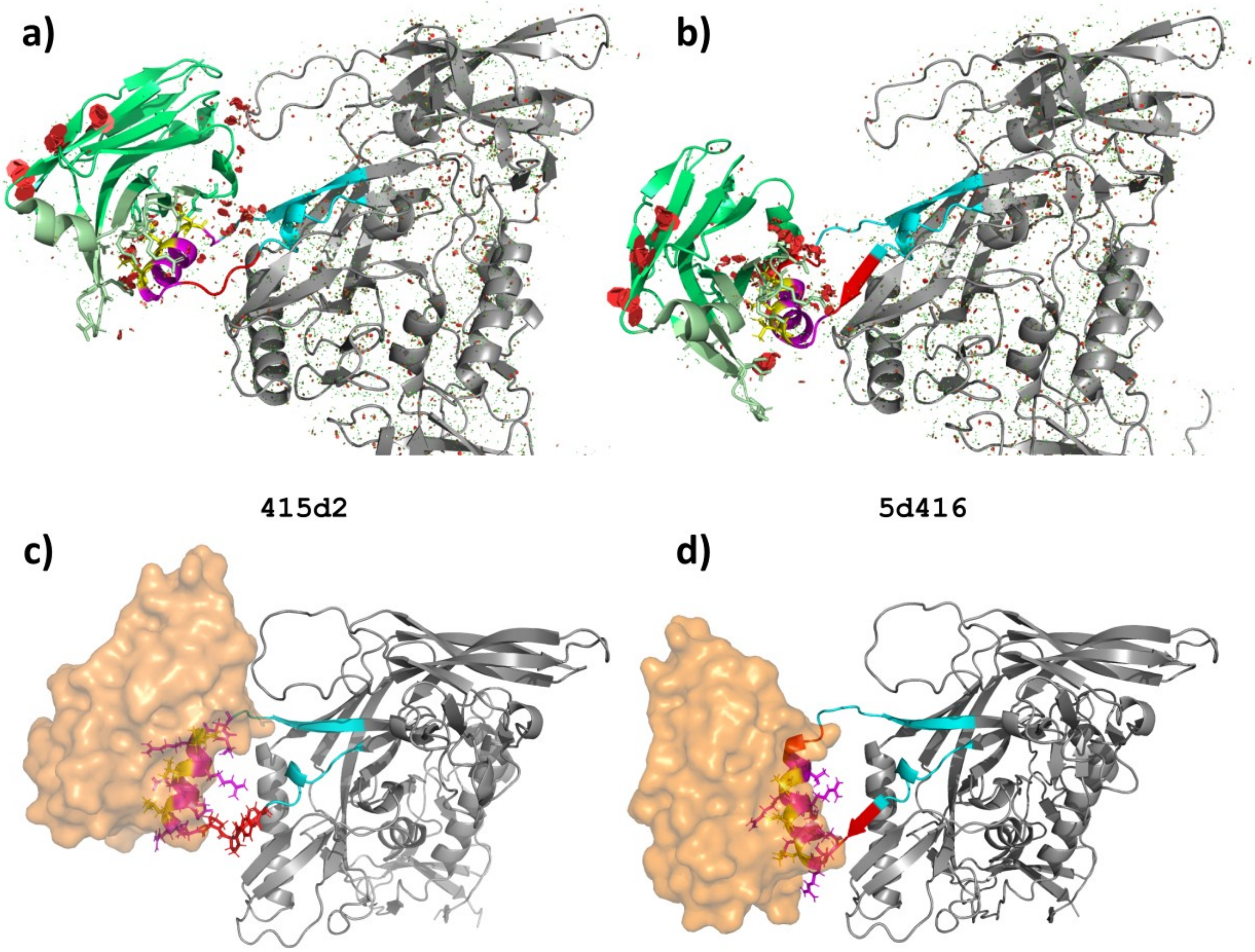


**Supplementary Figure 5. Visualizing nanobody alignment. (a, c)** JRFL V4-ALFA-tag UID 415d2 with the inserted tag aligned to the ALFA-tag from PDB ID 6IG2 to indicate where the NbALFA (shown in green cartoon or orange surface renderings) may bind. Steric clashes are indicated with red polygons. **(b, d)** JRFL V4-ALFA-tag UID 5d416, as in (a,c).

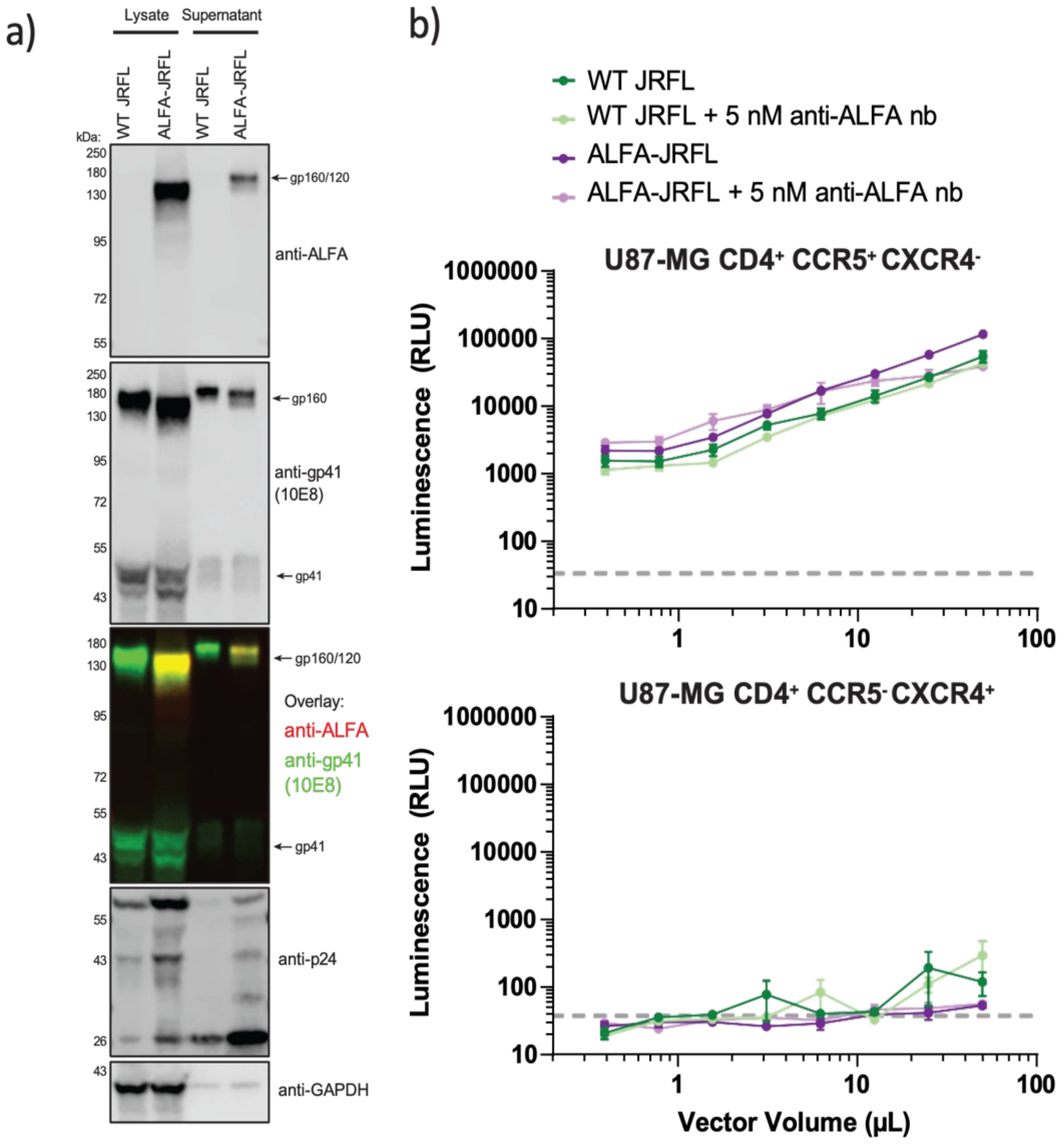


**Supplementary Figure 6. Validation of ALFA-Tagged HIV-1 JR-FL Env. (a)** Western blot of HEK293T lysates and supernatant transfected to produce HIV-1-based lentivirus pseudo typed with either wild-type or ALFA-tagged JR-FL (UID 415d2). **(b)** U87-MG CD4$^{+}$ cells expressing either CCR5 or CXCR4 were infected with a dilution series of WT or ALFA-tagged JR-FL pseudovirus, with or without a 30-minute pre-incubation of 5 nM NbALFA. Infection is measured as relative luminescence units.

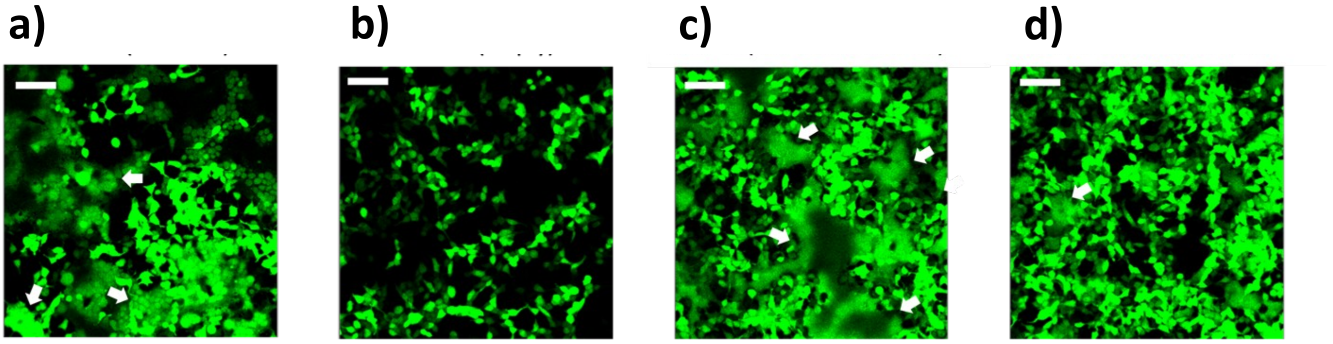


**Supplementary Figure 7. Cell-cell fusion assays for the JR-FL Env V4-ALFA-tag UID 415d2.** Permissive TZM-bl target cells were exposed to effector cells, HEK T293 cells co-expressing JR-FL Env: (a) wildtype, no V4 label, (b) empty vectors, (c) JR-FL V4-ALFA-tag (UID 415d12), and (d) JR-FL V-ALFA-tag UID 5d416. White arrows designate syncytia, i.e. cells that underwent fusion. Scale bar 10 μm.

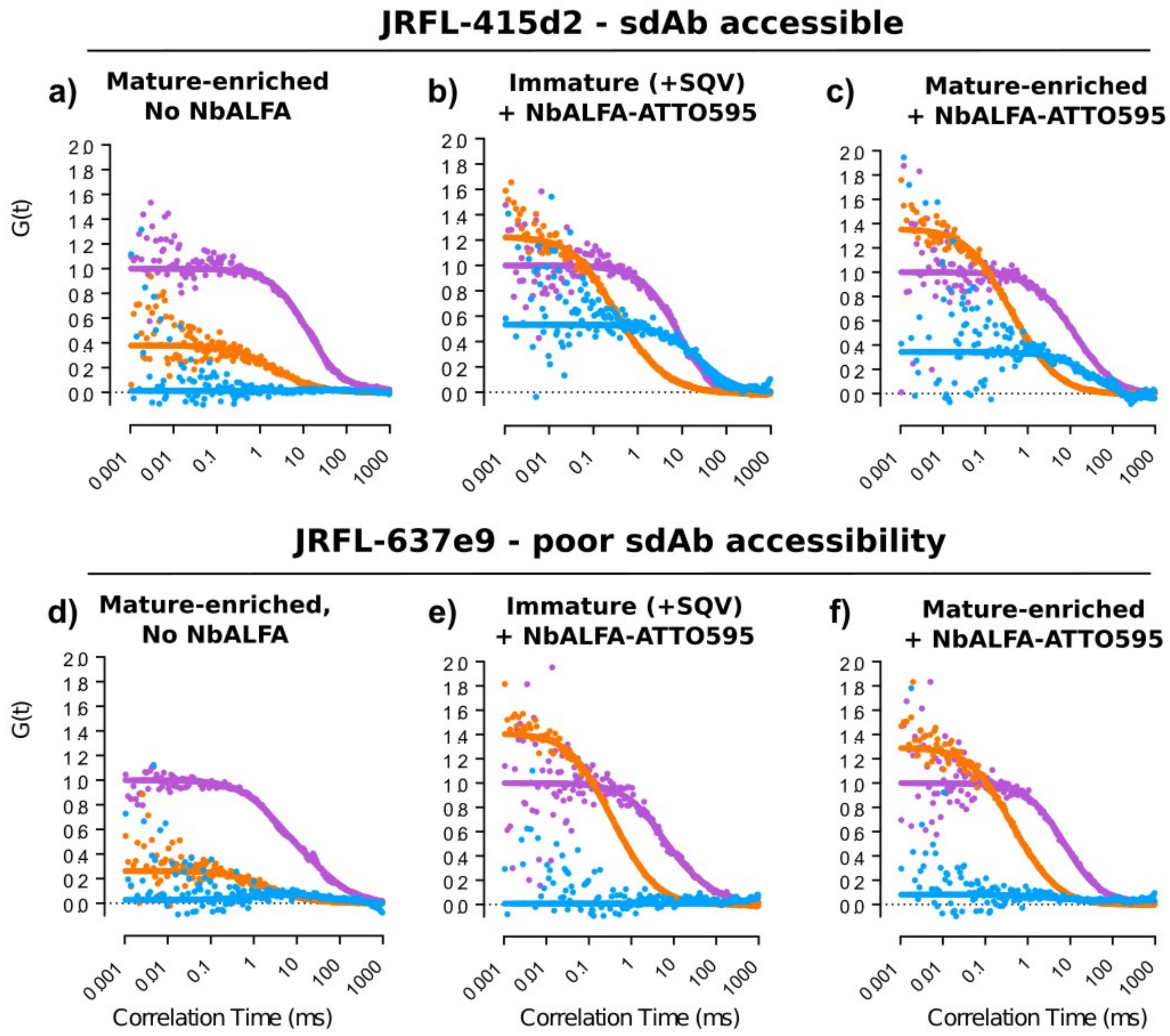


**Supplementary Figure 8. Fluorescence Cross-Correlation Spectroscopy (FCCS) demonstrates high specificity for nanobody engagement with JR-FL Env V4-ALFA-tag (UID 415d2) as indicated by Alphafold2.** Autocorrelation functions (ACF) for NbALFA-ATTO 594 (orange), Gag-EGFP (orange) and cross-correlation function (CCF, blue) for viral particles bearing either JR-FL Env V4-ALFAtag UID 415d2, bearing a competent tag structure **(a-c)** or UID 637e9, bearing an unfavourable tag structure **(d-f)**. ACF and CCF curves are shown for viral particles alone, i.e. without NbALFA **(a, d)**, immature (saquinivir treated) particles stained with NbALFA-ATTO 594 **(b, e)**, and mature (untreated) particles stained with NbALFA-ATTO 594 **(c, f)**. Cross-correlation between the virus (*via* Gag-EGFP) and the NbALFA diffusion is only indicated for the ALFA-tag insertion proposal which was expected to have an intact and likely accessible tag, i.e. UID 415d2.

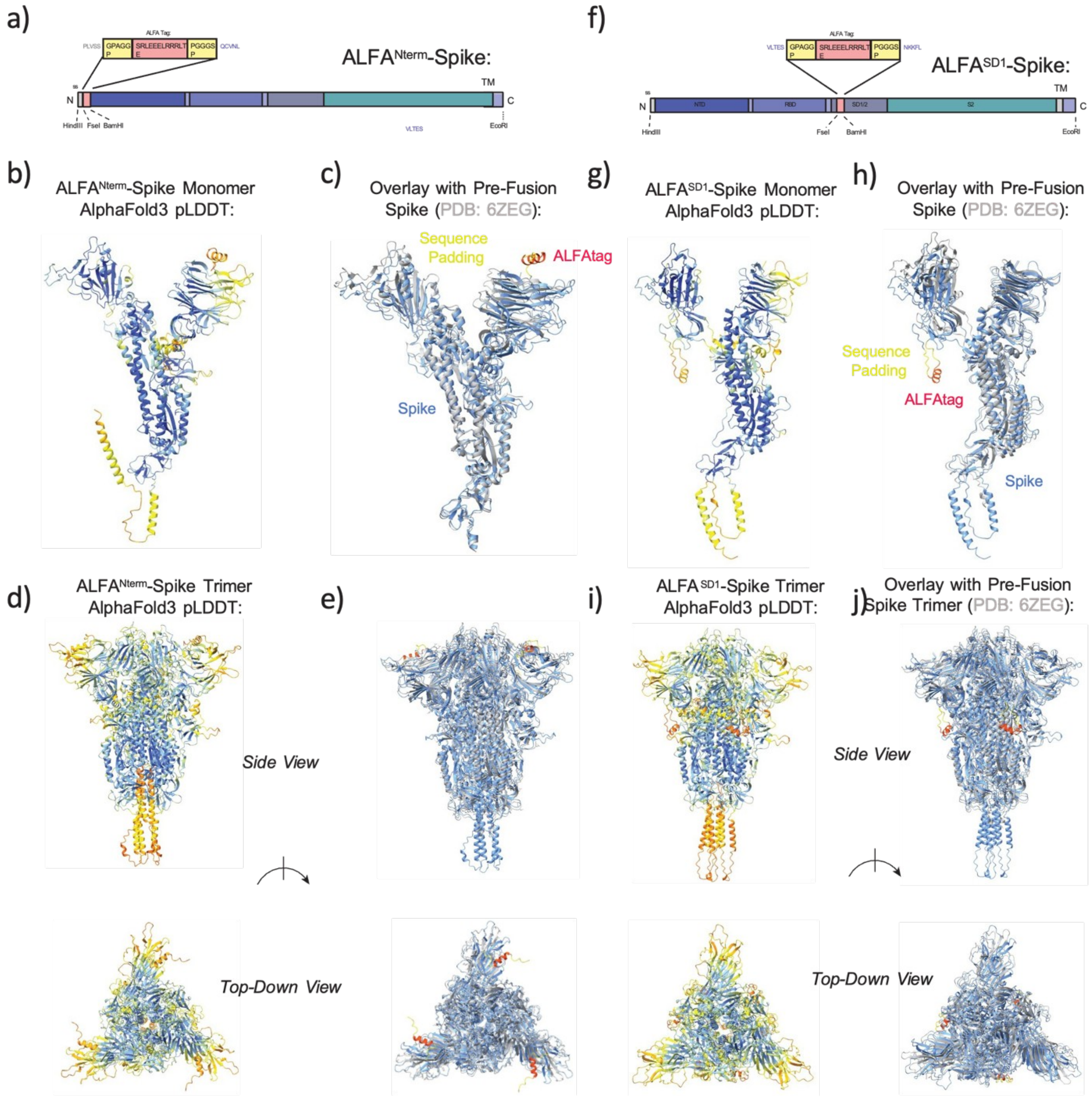


**Supplementary Figure 9. AlphaFold Representations of SARS-CoV-2 Spike with ALFA-tag insertions.** **(a)** Schematic of ALFA-tag insertion after the SARS-CoV-2 signal sequence to produce ALFA$^{Nterm}$-Spike. **(b)** AlphaFold3 model of ALFA$^{Nterm}$-Spike monomer, colored by pLDDT residue confidence score. **(c)** Overlay of ALFA$^{Nterm}$-Spike monomer with a monomer of the SARS-CoV-2 pre-fusion trimer cryo-electron microscopy structure (PDB ID 6ZEG). ALFA-tag and surrounding inserted residues are colored red and yellow, respectively. **(d)** AlphaFold 3 model of ALFA$^{Nterm}$-Spike trimer, colored by pLDDT residue confidence score. **(e)** Overlay of ALFA$^{Nterm}$-Spike trimer model with the SARS-CoV-2 trimer cyro-electron microscopy structure. **(f)** Schematic of ALFA-tag insertion in the unfolded SD1 domain of the SARS-CoV-2 Spike sequence to produce ALFA$^{SD1}$-Spike. **(g-j)** AlphaFold 3 models of the ALFA$^{SD1}$-Spike, colored by pLDDT or overlaid relative to PDB ID 6ZEG as in **(b-e)**.

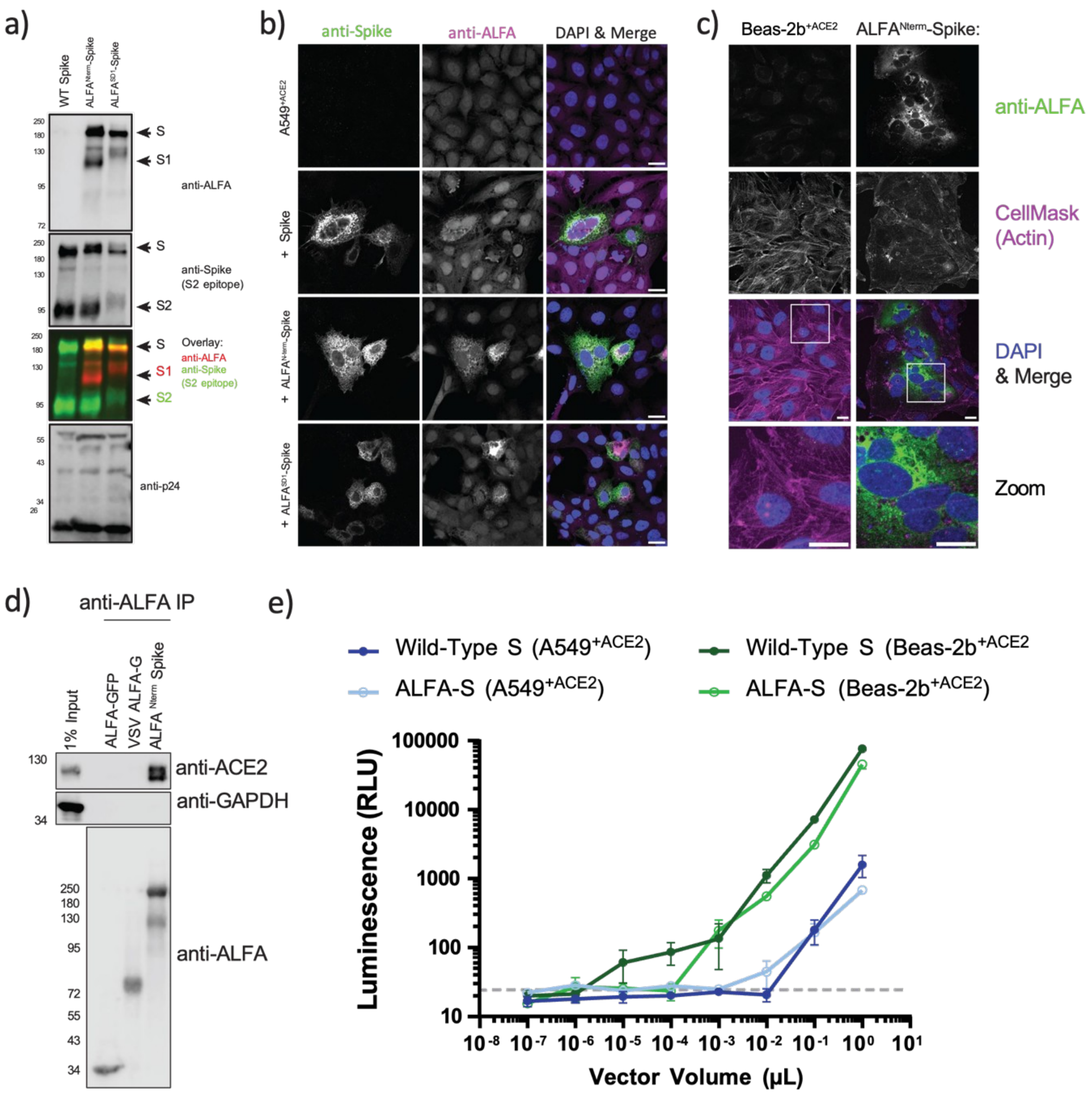


**Supplementary Figure 10. Functional Validation of SARS-CoV-2 ALFA-tagged Spikes**. **(a)** Western blot of concentrated supernatant from HEK293T cells producing lentiviral particles pseudo typed with wild-type, ALFA$^{Nterm}$-Spike or ALFA$^{SD1}$-Spike. The respective ALFA and S2 bands indicate productive cleavage at the S1/S2 cleavage site for wild-type and ALFA$^{Nterm}$-Spike. **(b)** Immunofluorescence microscopy of A549+ACE2 cells transfected with wild-type, ALFA$^{Nterm}$-Spike or ALFA$^{SD1}$-Spike. Wild-type and ALFA$^{Nterm}$-Spike facilitate cell-cell fusion events, whereas ALFA$^{SD1}$-Spike does not. **(c)** Representative immunofluorescence micrograph of BEAS-2B$^{+ACE2}$ cells transfected with ALFA$^{Nterm}$-Spike and resulting syncytia formation. **(d)** ALFA$^{Nterm}$-Spike retains ACE2-binding capacity. HEK293T$^{+ACE2}$ cells were transfected with ALFA-GFP, VSV-ALFA G or SARS-CoV-2 ALFA$^{Nterm}$-Spike. Cells were lysed, then incubated with ALFA Selector beads. Eluates were immunoblotted for ACE2, or GAPDH as a negative control. **(e)** Luminescence values for A549$^{+ACE2}$ and BEAS-2B$^{+ACE2}$ cells incubated with serial dilutions of lentiviruses pseudotyped with wild-type Spike or ALFA$^{Nterm}$-Spike.

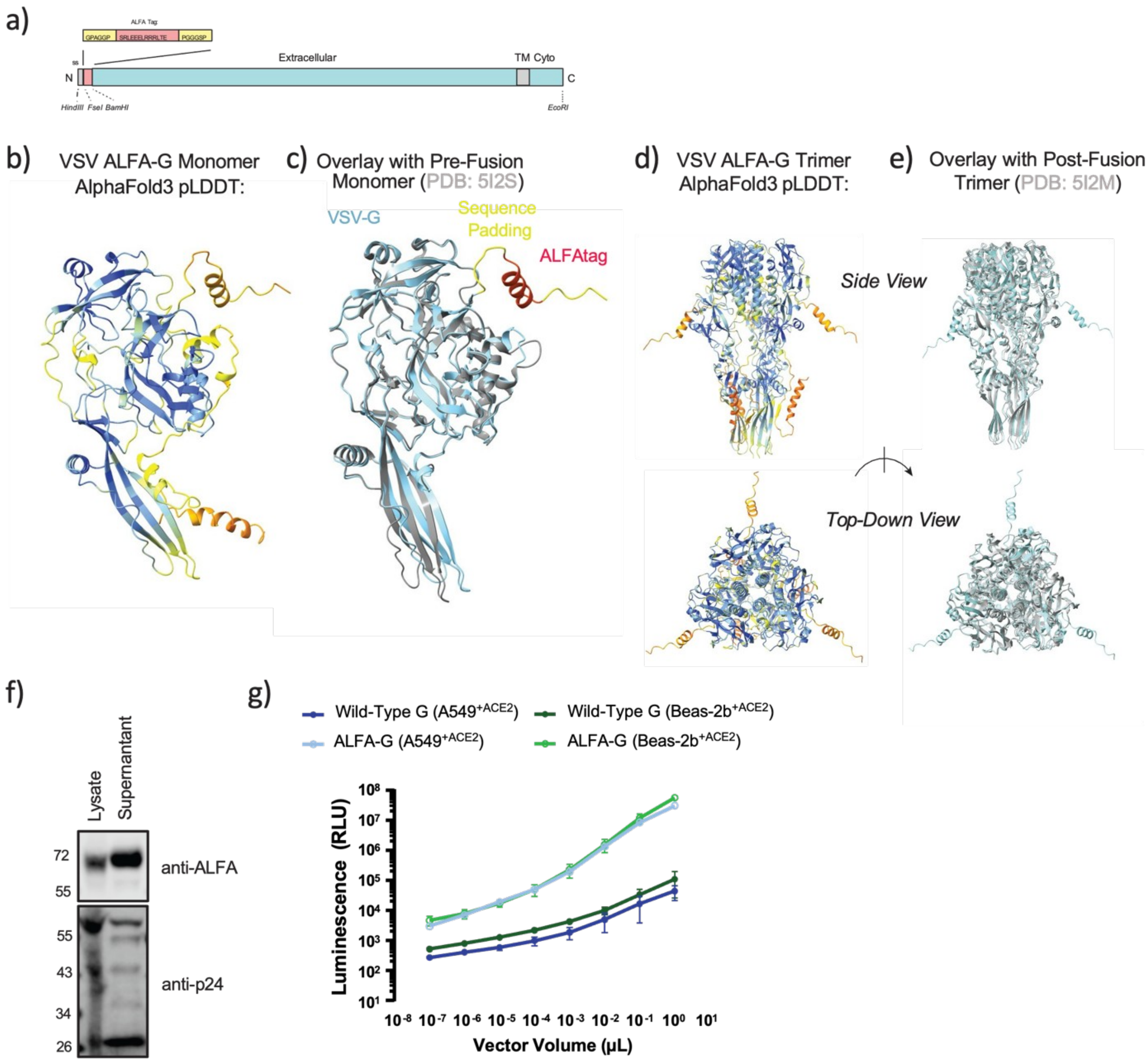


**Supplementary Figure 11. Design, Modeling, and Validation of VSV ALFA-G**. **(a)** Schematic of ALFA-tag insertion after a signal peptide sequence to produce an N-terminally tagged ALFA-G protein. **(b)** AlphaFold2 model of an ALFA-G monomer, colored by pLDDT residue confidence score. **(c)** Overlay of ALFA-G monomer with a pre-fusion monomer structure (PDB ID 5I2S). ALFA-tag and surrounding inserted residues are colored red and yellow, respectively. **(d)** AlphaFold2 model of ALFA-G, colored by pLDDT residue confidence score. **(e)** Overlay of ALFA-G trimer model with the VSV-G trimer cyro-electron microscopy structure (PDB: 5I2M). **(f)** Western blot of cell lysate and concentrated supernatant from HEK293T cells producing lentiviral particles pseudotyped with ALFA-G. **(g)** Luminescence values for A549$^{+ACE2}$ and BEAS-2B$^{+ACE2}$ cells incubated with serial dilutions of lentiviruses pseudotyped with ALFA-G.

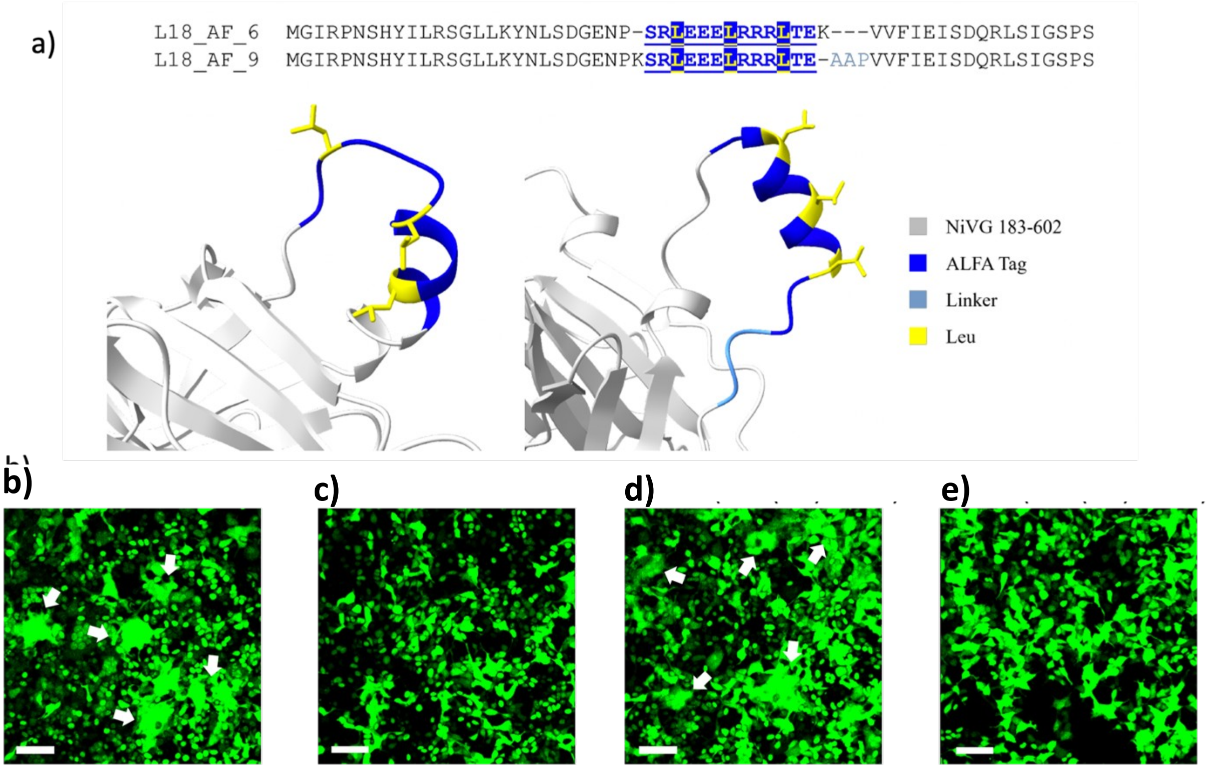


**Supplementary Figure 12. ALFA-tag Modelling for Nipah G protein. (a)** Sequences of two variants with ALFA-tag and linker amino acids inserted into loop 18, GENPK. *Left:* Variant L18_AF_3, predicted with non-alpha helical ALFA-tag folding. *Right:* Variant L18_9, predicted with alpha-helical ALFA-tag after insertion of a linker sequence. **(b)** Cell-cell fusion experiments where target (HEK T293 cells) were exposed to effector cells, Lenti-X 293TΔEFNB2 cells co-expressing wild type Nipah G and Nipah F proteins, **(c)** empty vectors, **(d)** Nipah G (AF5 tag variant) + Nipah F which presented syncytia, white arrows, and behaves as the wild type and **(e)** Nipah G (AF4 tag variant) + Nipah F which efficiency is ostensibly diminished compared to the wild type example (b) and the AF5 tag variant (d). White arrows designate cells that underwent fusion. Scale bar 10 µm.

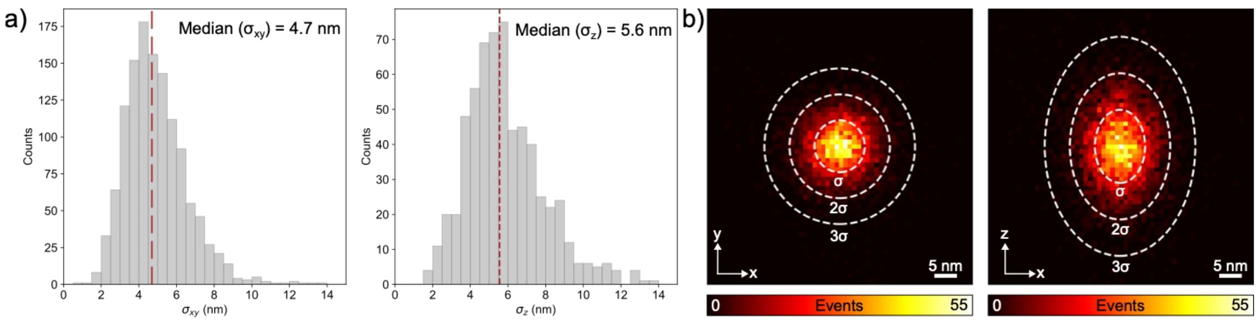


**Supplementary Figure 13. Localization precision in 3D MINFLUX imaging. (a)** Lateral and axial precision of localizations for all single molecule fluorescence events for same field of view shown in Fig. 2b. **(b)** Histogram of the distance of a localization to the mean position of a single fluorophore. The ellipses are displayed with semi-axes of σ, 2σ, and 3σ in length, with σ the precision obtained from a combined analysis of the statistical localization spread (standard deviation) in xy, and xz.

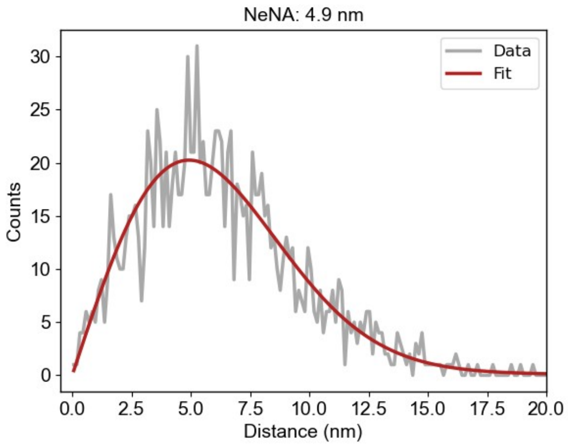


**Supplementary Figure 14**. Overall localization precision of all super-resolution DNA-PAINT images of single mature virus-like particles based on Nearest- Neighbor analysis (NeNa).

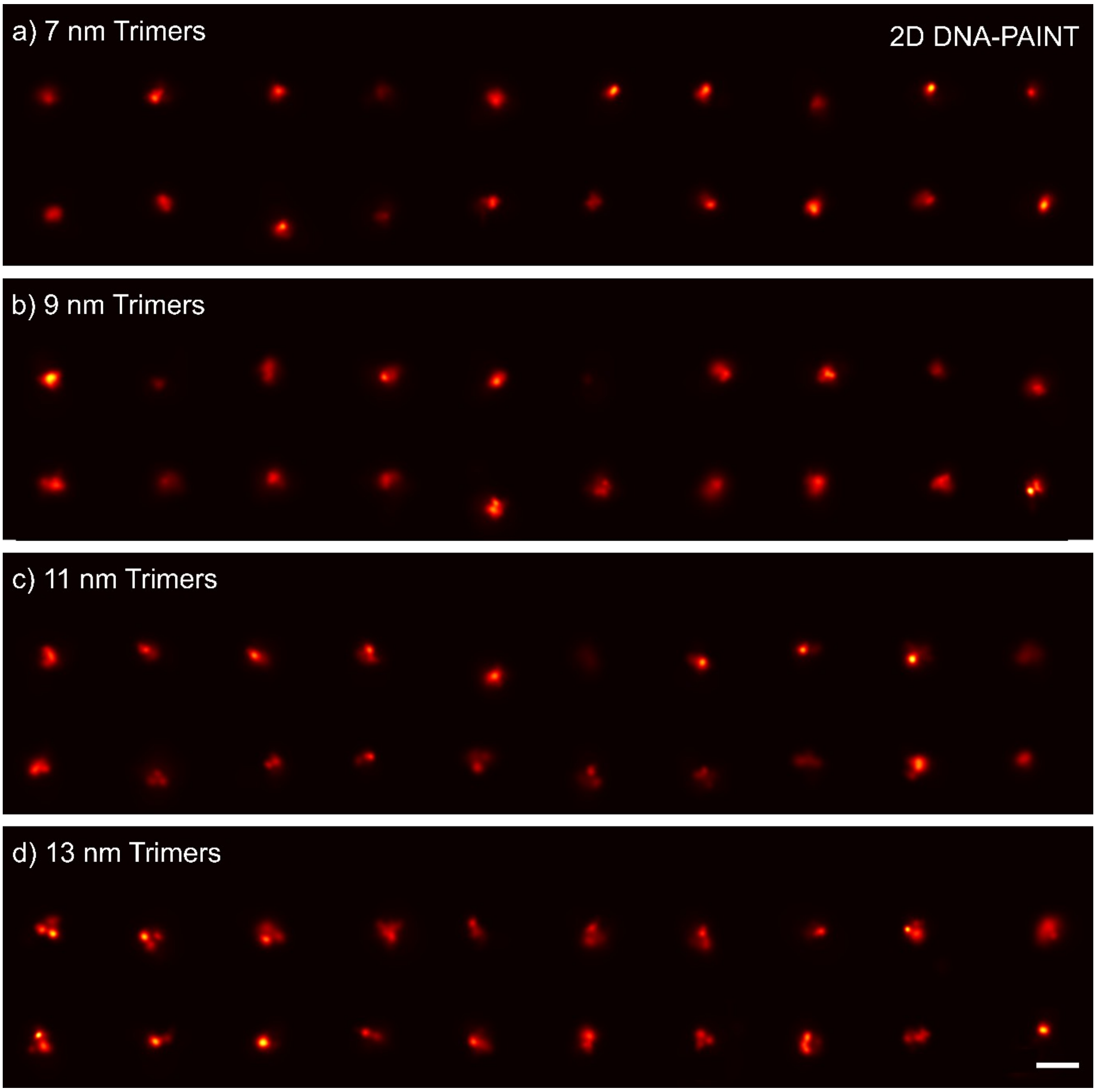


**Supplementary Figure 15: 2D DNA-PAINT simulations for equilateral trimers with different distances (a)** 20 trimers of 7 nm distances between them and 80% labelling efficiency. **(b)** 20 trimers of 9 nm distances between them and 80% labelling efficiency. **(c)** 20 trimers of 11 nm distances between them and 80% labelling efficiency. **(d)** 20 trimers of 13 nm distances between them and 80% labelling efficiency. All simulations were performed using the same parameters as the experimental conditions, including kinetics and signal-to-background ratio. Scale bar: 50 nm.
